# Single cell transcriptome profiling of the human developing spinal cord reveals a conserved genetic programme with human specific features

**DOI:** 10.1101/2021.04.12.439474

**Authors:** Teresa Rayon, Rory J. Maizels, Christopher Barrington, James Briscoe

## Abstract

The spinal cord receives input from peripheral sensory neurons and controls motor output by regulating muscle innervating motor neurons. These functions are carried out by neural circuits comprising molecularly and physiologically distinct neuronal subtypes that are generated in a characteristic spatial-temporal arrangement from progenitors in the embryonic neural tube. The systematic mapping of gene expression in mouse embryos has provided insight into the diversity and complexity of cells in the neural tube. For human embryos, however, less information has been available. To address this, we used single cell mRNA sequencing to profile cervical and thoracic regions in four human embryos of Carnegie Stages (CS) CS12, CS14, CS17 and CS19 from Gestational Weeks (W) 4-7. In total we recovered the transcriptomes of 71,219 cells. Analysis of progenitor and neuronal populations from the neural tube, as well as cells of the peripheral nervous system, in dorsal root ganglia adjacent to the neural tube, identified dozens of distinct cell types and facilitated the reconstruction of the differentiation pathways of specific neuronal subtypes. Comparison with existing mouse datasets revealed the overall similarity of mouse and human neural tube development while highlighting specific features that differed between species. These data provide a catalogue of gene expression and cell type identity in the developing neural tube that will support future studies of sensory and motor control systems and can be explored at https://shiny.crick.ac.uk/scviewer/neuraltube/.

## Introduction

The spinal cord receives and processes information from sensory neurons in the peripheral nervous system (PNS) and controls muscle movement by coordinating the activity of motor neurons (MNs). The neural circuits that perform these tasks comprise molecularly and physiologically distinct classes of neurons that are generated in a stereotypic spatial and temporal order from proliferating progenitors in the embryonic neural tube and neural crest. The spatiotemporal arrangement of progenitors and the identity of the neurons they generate is determined by gene regulatory networks (GRNs) controlled by extrinsic signals (Delás and Briscoe, 2020; Sagner and Briscoe, 2019). The composition and operation of these GRNs have been studied in model organisms, such as mouse, chick and zebrafish and recent work has included the systematic profiling of single cells from the mouse neural tube and neural crest (Delile et al., 2019; Ray et al., 2018; Rosenberg et al., 2018; Sharma et al., 2020; Soldatov et al., 2019). This has provided catalogues of gene expression, revealed complexity, and allowed the detailed molecular classification of multiple cell types.

Although recent attention has focused on profiling the transcriptomes of cells in several regions of the developing human brain (Eze et al., 2021), the embryonic human spinal cord and peripheral nervous system have been less well described. The available molecular characterization has shown that the identity and overall organization of progenitors and neurons is, in the main, similar between humans and other vertebrates (Dady et al., 2021; Marklund et al., 2014; Rayon et al., 2020). Nevertheless, the extent of the similarity in the molecular composition of cells in mouse and human neural tubes has not been determined. Notably, several features of human neural tube development have been reported to differ from other species. For example, a subset of neural progenitors in the spinal cord coexpress the transcription factors OLIG2 and NKX2.2 in human embryos. These are relatively rare at the equivalent developmental stages in the mouse and chick (Marklund et al., 2014).

Systematic profiling of cell identity in the developing nervous system of human embryos would offer a clearer picture of the developing human neural tube and allow comparisons with other species. To this end we profiled gene expression in single-cell transcriptomes in samples of dissected trunks from human embryos from Gestational Week (W) 4 to 7. We recovered a total of 71,219 cells, of which 43,485 were neural, from 4 embryos and used these to generate a single-cell atlas of the developing human spinal cord and dorsal root ganglia. Comparison with equivalent staged mouse embryos identified similarities and differences. To allow the data to be explored further we developed an online resource that is available at https://shiny.crick.ac.uk/scviewer/neuraltube/.

## Results and Discussion

### Identification of cell types in developing human embryos

To identify the cell types in the developing human spinal cord, we performed single-cell RNA sequencing (scRNA-seq) using the droplet-based 10X Chromium platform. We microdissected cervical-thoracic regions from shoulder to hip level of single embryos during the first trimester of human development, spanning four weeks of development from W4 to 7, corresponding to Carnegie Stages (CS) 12, 14, 17 and 19. The samples from CS12, CS14 and CS19 were processed on the same day as the termination of pregnancy, whereas CS17 was processed for sequencing after a 24h delay. We dissected CS14, CS17 and CS19 samples into anterior and posterior regions and sequenced these separately. Since the CS12 sample was smaller in size, to minimize cell loss we avoided further regional microdissection and sequenced cells of the entire trunk element. In addition, to compensate for the increased cell numbers of the older embryos, two replicates per timepoint were generated for the CS17 and the CS19 samples. In total, 120,620 droplets were annotated as cells after sequencing. To each sample, we applied similar quality filters to those we established for the developing mouse spinal cord (Delile et al., 2019). This resulted in a dataset of 71,219 cells: 11,963 from CS12; 11,525 from CS14; 28,800 from CS17; and 18,931 from CS19 (FIG S1A-D). More cells were detected in the CS17 sample compared to the other samples. This was likely due to the increased ambient RNA as indicated by the increased dispersion in the proportion of mitochondrial UMIs. The total number of detected genes (median ∼3,000 per cell) and unique molecular identifiers (UMIs) in cells were similar in all samples analysed (FIG S1A,B), and were similar to the mouse single cell dataset (Delile et al., 2019). Expression of the male-specific *SRY* gene (Berta et al., 1990; Gubbay et al., 1990; Sinclair et al., 1990) was detected in CS12 and CS17 embryos, indicating these samples were male, whilst the lack of SRY expression in the CS14 and CS19 samples indicated that these were female (FIG S1F).

In a first step, we visualized each sample separately using Uniform Manifold Approximation and Projection (UMAP). To allocate cell types within this embedding, we calculated a gene module score (see Methods) that included the marker genes used to characterize tissue types in mouse (Delile et al., 2019). This identified progenitors and neurons of the spinal cord and dorsal root ganglion, mesoderm, hematopoietic and skin cells (FIG 1A). We labeled as “other” cells with poorly resolved transcriptomes and cells derived from other tissues of the embryo. Each of the samples displayed a similar composition of cell types (FIG 1A) but with obvious differences in the proportions at different timepoints (FIG 1B). More neurons were contained in the CS19 dataset, consistent with the progression of neurogenesis in the spinal cord over time. By contrast, mesodermal cells were enriched in the early stages (CS12 and CS14) compared to the late timepoints (FIG1B). This is due to the proximity of the mesoderm and neural tube at early stages, and the greater difficulty of cleanly dissecting smaller embryos.

**Figure 1.**
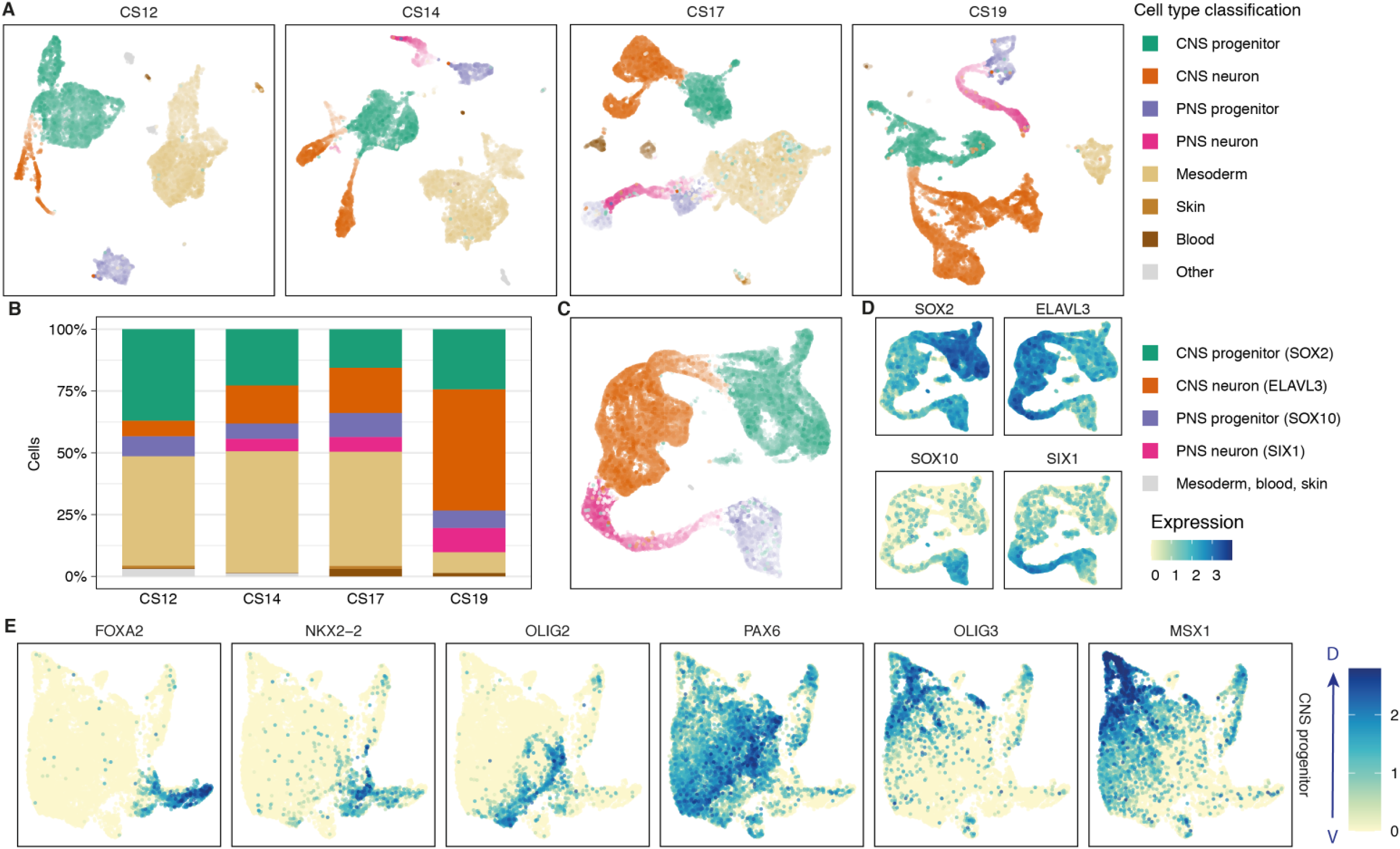
Single cell RNA-seq from the human developing spinal cord. **(A)** UMAP of CS12, CS14, CS17 and CS19 samples annotated by gene module score. **(B)** Stacked bar chart indicating the cell type composition per timepoint. **(C)** UMAP of the combined human neural dataset colored by gene module score. Colour and brightness of cells reflects the representative gene module (n=43,485). **(D)** Feature plots of markers of neural progenitors (SOX2), neurons (ELAVL3) and neural crest progenitors (SOX10) and neurons (SIX1). **(E)** Feature plots of the spinal cord progenitor UMAP showing the patterned expression of the indicated DV patterning markers.

To generate a spatiotemporal gene expression atlas of developing neural cells, we selected the neural cell clusters from each stage and combined these into a single dataset for further analysis. The combined human UMAP of the neural dataset contained a total of 43,485 cells and allowed us to distinguish progenitors and neurons of both the neural tube and dorsal root ganglia. Cells from different samples, whether of the same or different sex, did not form distinct clusters within the embedding, which suggested minimal batch effects after data processing (FIG S1G). Expression of the progenitor marker *SOX2* and the neuronal marker *ELAVL3* allowed the identification of progenitors and neurons within the map. Likewise, the expression of the neural crest markers *SOX10* and *SIX1* allowed us to discriminate progenitors and neurons of neural crest origin (6,747 cells) versus spinal cord (32,928 cells) (FIG 1C). Within the UMAP, neurons of both the central and peripheral nervous system projected to adjacent clusters, pointing to a convergence in the gene expression programmes of central and peripheral neurons (FIG 1C).

### Classification of central nervous cells

Computationally isolating spinal cord progenitor cells from all four timepoints arranged cells in the UMAP in a way that resembled the dorsoventral (DV) patterning of the developing neural tube (FIG 1E). Nevertheless, attempts to use standard unsupervised methods to cluster progenitors were unsuccessful at classifying cells into the known dorsoventral domains. Similar approaches with neurons were also unsuccessful. This suggested that, similar to mouse, the complexity and combinatorial expression profile of genes in both progenitors and neurons preclude the use of unsupervised approaches.

In order to classify subtype identity of neural cells, we adapted the ‘knowledge matrix’ we had previously constructed, using available molecular characterization of the vertebrate developing neural tube (Delile et al., 2019) (Table S2). This represents an inventory in which each progenitor type and neuronal class is defined by the presence or absence of a characteristic set of marker genes (FIG 2B). We binarized the expression levels of the marker gene set in the human transcriptome data and assigned each cell an identity from the knowledge matrix. This allowed the categorization of cell types into specific progenitor and neuronal domains (FIG 2A-C). Plotting the scaled mean expression of the genes used to assign cell identities revealed the expected conserved patterns of progenitor- and neuron-specific gene expression along the dorsoventral axis of the spinal cord (FIG 2A-C). Overall, we observed similar identities and organization of progenitors and neurons in mouse and human (FIG S2A,B) with a mean scaled expression for most genes comparable between mouse and human. To compare the transcriptional similarity between progenitors and neurons of mouse and human, we used expression levels of homologous genes in each neural cell type in mouse and human and estimated the Pearson correlation coefficient for each pair of cell types. We measured the correlation using a set of transcription factors (see methods). Mouse and human cells assigned to the same cell type had the highest correlation (FIG 2D). The similarity of progenitor cells was higher between adjacent dorsoventral domains. By contrast, neuronal cell types clustered according to their excitatory or inhibitory function (Sagner and Briscoe, 2019) (FIG 2D).

**Figure 2.**
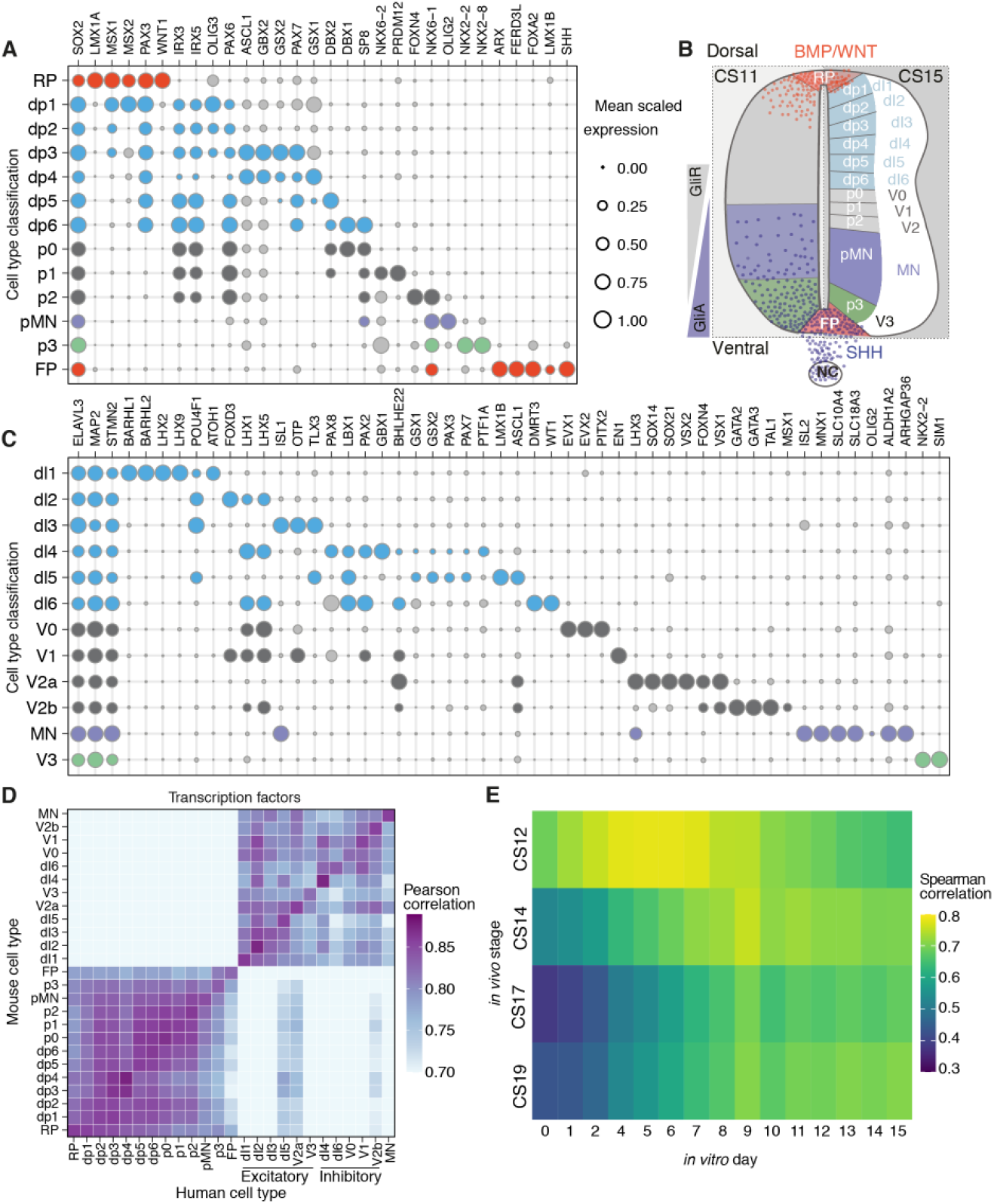
Classification of dorsoventral progenitors and neuronal classes in human. **(A)** Bubble plots indicating the expression of markers used to identify dorsoventral (DV) domains in human progenitors. **(B)** Diagram of the DV domains in the human developing spinal cord highlighting the opposing gradients of SHH and WNT/BMP and the patterning of the 11 DV domains, the floor plate and roof plate. **(C)** Bubble plots depicting the 11 neuronal classes in human. In (A) and (C), genes chosen for cell assignment are coloured; grey circles correspond to markers not used for the selection of a specific population. Circle size indicates mean scaled gene expression levels. **(D)** Heatmap of the pairwise Pearson correlation coefficients of gene expression in mouse (vertical) and human (horizontal) progenitor and neural cells. **(E)** Heatmap of Spearman correlation coefficients per timepoint of the averaged gene expression of ventral progenitors and neurons in the human spinal cord *in vivo* (vertical) and *in vitro* (horizontal).

The availability of the human neural tube data allowed us to make comparisons with the differentiation of MNs *in vitro* from human embryonic stem cells. We selected highly expressed genes that varied during neurogenic differentiation from daily bulk RNAseq samples of an *in vitro* differentiation of human MNs (Rayon et al., 2020). We then generated “pseudo bulk RNA” samples from the human *in vivo* dataset by averaging gene expression of progenitors and neurons assigned to ventral domains (p3, pMN, p2, p1 and p0) at each timepoint. Pair wise comparison of *in vitro* and *in vivo* timepoints indicated that the CS12 sample had the highest correlation to in vitro Day 5 (FIG 2E). CS14 was most similar to *in vitro* Day 9. Thus, the four-day *in vitro* differentiation from Day 5 to Day 9 was similar to the developmental progression from CS12 to CS14, corresponding to gestational days 26-30 and days 31-35, respectively. By contrast, CS17 and CS19 stages, which correspond to gestational days 42-44 and day 48-51 of human development, showed higher correlations with later *in vitro* timepoints albeit there was less similarity at these stages. At least in part this could be because *in vitro* differentiations become progressively more asynchronous at later time points.

### Specific features of gene expression in the human spinal cord

Although the transcriptomes of mouse and human neural cells were broadly similar, specific differences between mouse and human were apparent. *PAX7* expression was observed in dorsal progenitors of both mouse and human, but in addition *PAX7* expression was observed in floor plate cells of CS12, CS14 and CS17 human embryos. However, it was absent from mouse floor plate cells at all timepoints (FIG 2A, S2A). This is consistent with immunohistochemical analysis of PAX7 in the human spinal cord from CS12 to CS15 (Betters et al., 2010; Dady et al., 2021). Inspection of genes correlating with *PAX7* in human FP cells highlighted several, including *CDH7, TXLNB, CDHR3, PIFO* and *RRAD*, that were expressed in human FP cells but not in mouse FP cells (FIG 3A-C). Whether *PAX7* directly regulates these or other genes in human FP cells will require functional experiments.

**Figure 3.**
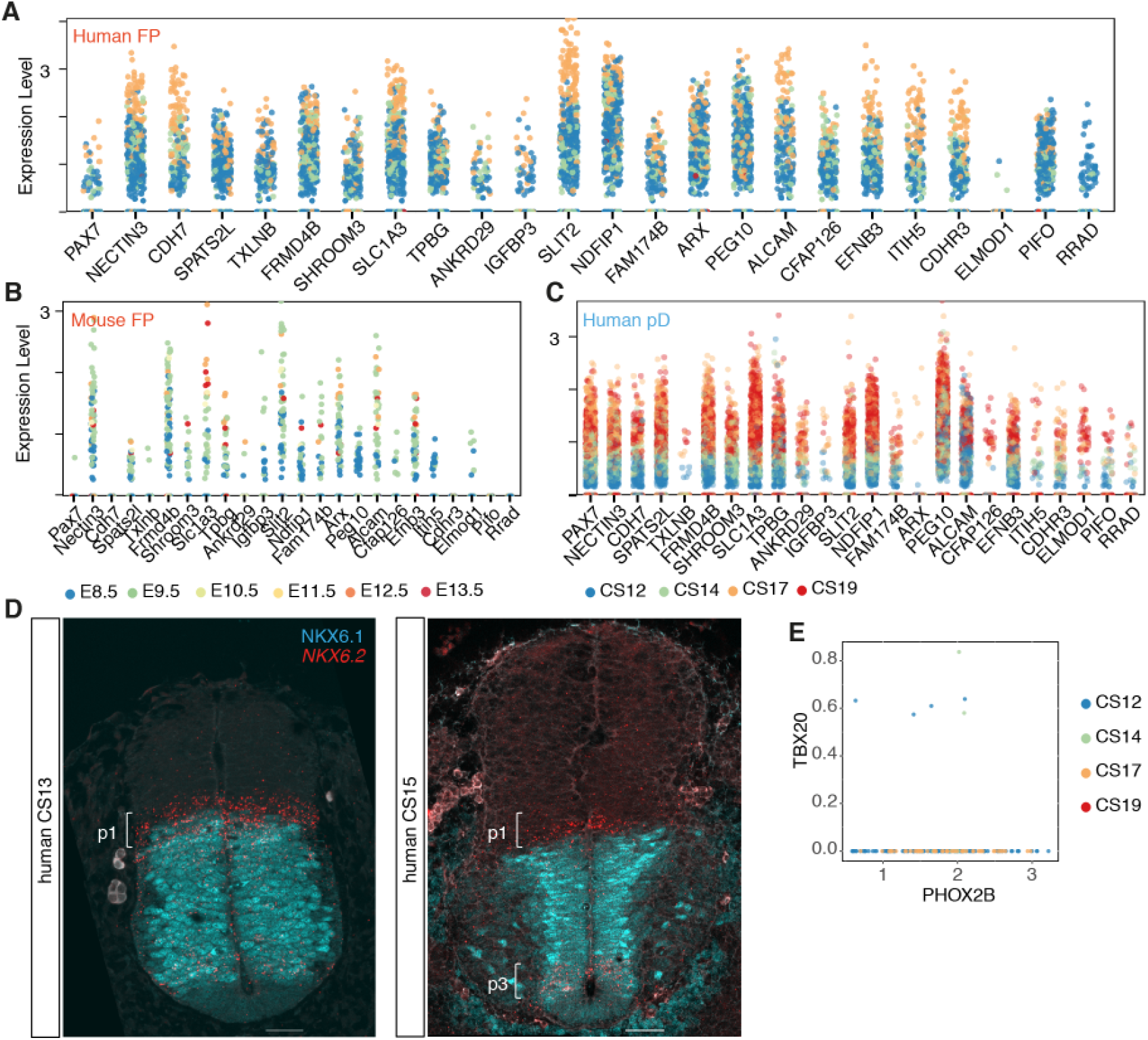
Human-specific features of neural progenitors and visceral motor neurons. **(A-C)** Genes most correlated with *PAX7* in (D) human FP, (E) mouse FP, and (F) in human pD (dp1-dp6). **(D)** NKX6.1(cyan) and *NKX6*.2 (red) expression in transverse sections of the human neural tube at shoulder levels in CS13 and CS15 embryos. Scale bars, 50μm. **(E)** Scatter plot of the expression of *PHOX2B* and *TBX20* in human (n=170 cells).

*NKX6*.*2* expression in human was detected in p1, p2, pMN, p3 and FP progenitors (FIG 2A). By contrast, mouse *Nkx6*.*2* defines p1 cells but is largely absent from other ventral progenitor domains (FIG S2A, S2C) (Vallstedt et al., 2001). The expression of *NKX6*.*2* in the p1 domain of the human neural tube was maintained across the timepoints analysed but decreased over time in the pMN and p3 domains (FIG 3D, S3B,C). The broad ventral expression profile of *NKX6*.*2* in the human neural progenitors resembled the broad expression of *Nkx6*.*2* in chick embryos and the transiently wider domain of expression of NKX6.2 at early development stages in the mouse (Vallstedt et al., 2001). This suggests that a heterochronic shift in the timing of *NKX6*.2 downregulation could account for the difference in expression between mouse and human.

Other notable differences between human and mouse included the apparent scarce expression of *TBX20* in human visceral MNs. These MNs, present in the hindbrain and cervical spinal cord, are characterized by the expression of *Isl1, Phox2b, Tbx2, Tbx3* and *Tbx20* in mouse (Pontecorvi et al., 2008). Analysis of *PHOX2B* expressing neurons in the human dataset revealed that although these cells expressed *TBX2* and *TBX3* there was little or no expression of *TBX20* (FIG 3E, S3D).

### Expression of primate-specific genes

The expression of genes specific to primates has received attention because of their potential role in species-specific aspects of human brain development, particularly in the neocortex (Fiddes et al., 2018; Florio et al., 2015; Heide et al., 2020; Liu et al., 2017; Suzuki et al., 2018). To investigate the expression of primate-specific genes in the more evolutionarily conserved spinal cord, we selected a list of 51 primate-specific protein-coding genes that have been shown to be enriched in human cerebral neural precursors (Florio et al., 2018; Liu et al., 2017) and assessed their expression. First, we examined the tissue specificity of expression. *TMEM133* and *ZNF788* were not detectable in the dataset, suggesting they might be brain specific. Of the remaining 49 genes, 87% (43/49) were expressed broadly in more than one tissue (FIG 4A), suggesting these were not specific to the nervous system. These included genes that have been shown to function in cortical expansion or folding in human (*TMEM14B, ARHGAP11B, NOTCH2NL*) (Fiddes et al., 2018; Florio et al., 2015; Heide et al., 2020; Liu et al., 2017; Suzuki et al., 2018). We examined more closely genes expressed at an average level of 0.10 transcripts/cell or greater in the CNS or PNS (9 genes out of 43; 20%). The only gene expressed above this threshold with documented functions in brain development was *TMEM14B* (FIG 4A). *TMEM14B* along with the other 8 genes showed widespread expression in progenitors and neurons, with a higher percentage of progenitors expressing these genes (FIG 4B,C). *CCDC47B, CBWD2, DHRS4* and *ZNF90* were somewhat more expressed in populations of CNS neurons compared to progenitors (FIG 4B,C). Thus, expression of primate-specific genes implicated in developmental and neurogenic functions can be detected in the spinal cord and other tissues (FIG S4A,B). The function of most of these genes remains unknown (Florio et al., 2018) and further analysis will be necessary.

**Figure 4.**
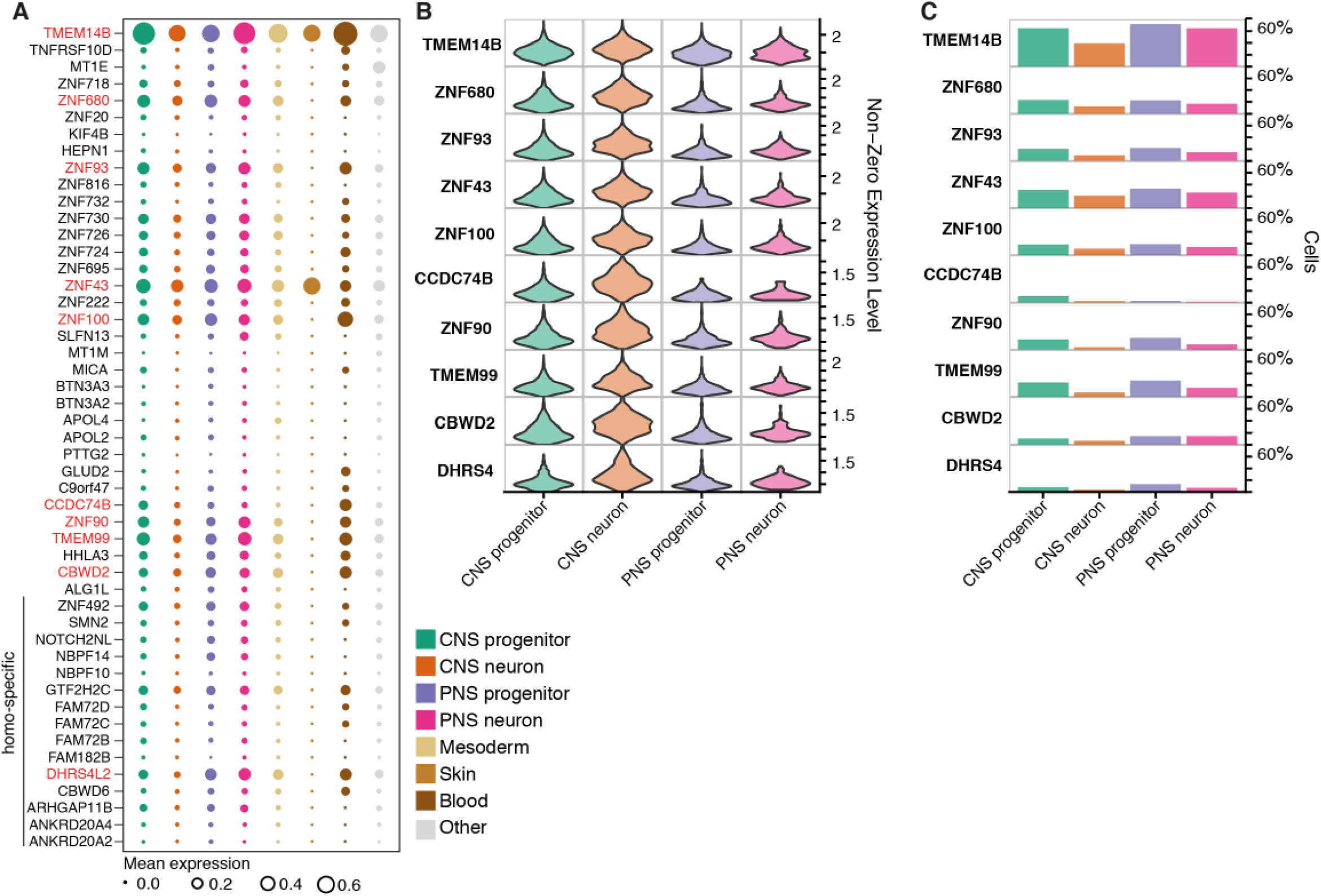
Expression of primate-specific genes. **(A)** Bubble plots indicating the expression of primate specific genes in different cell types. Genes expressed at an average level of 0.10 transcripts/cell or higher in any cell type are highlighted in red. Circle size indicates mean scaled gene expression levels. **(B)** Violin plots for the expression of genes in neural cell types. Cells with null values for the genes were not included. **(C)** Proportion of cells within each cell type that express the indicated primate-specific genes.

### Classification of peripheral nervous system cells

The peripheral nervous system (PNS) is derived from neural crest cells and emerges in trunk regions of human embryos around CS11-12 (O’Rahilly and Müller, 2007). This allowed us to follow the differentiation dynamics of neural crest cells to PNS neurons during human embryogenesis. To classify subtype identity of neural crest cells and investigate sensory neurogenesis in human, we mapped cells using markers of the PNS that had previously been defined in other species (Chiu et al., 2014; Hockley et al., 2019; Li et al., 2016; Usoskin et al., 2015; Vermeiren et al., 2020; Zeisel et al., 2018). We established a classification scheme to subdivide cells into the three cell types characteristic of the E11.5 mouse trunk dorsal root ganglia (Soldatov et al., 2019): (i) progenitors, marked by the expression of *SOX10* alone or in combination with *SOX2*; (ii) sensory neuron precursors, marked by the expression of *NEUROG1, NEUROG2* and *NEUROD1*; and (iii) postmitotic sensory neurons expressing *ELAVL3* (FIG 5A), as the expression of the broad mouse somatosensory neuron marker advillin (AVIL) was limited to a small subset of cells in the human dataset (FIG 5A).

**Figure 5.**
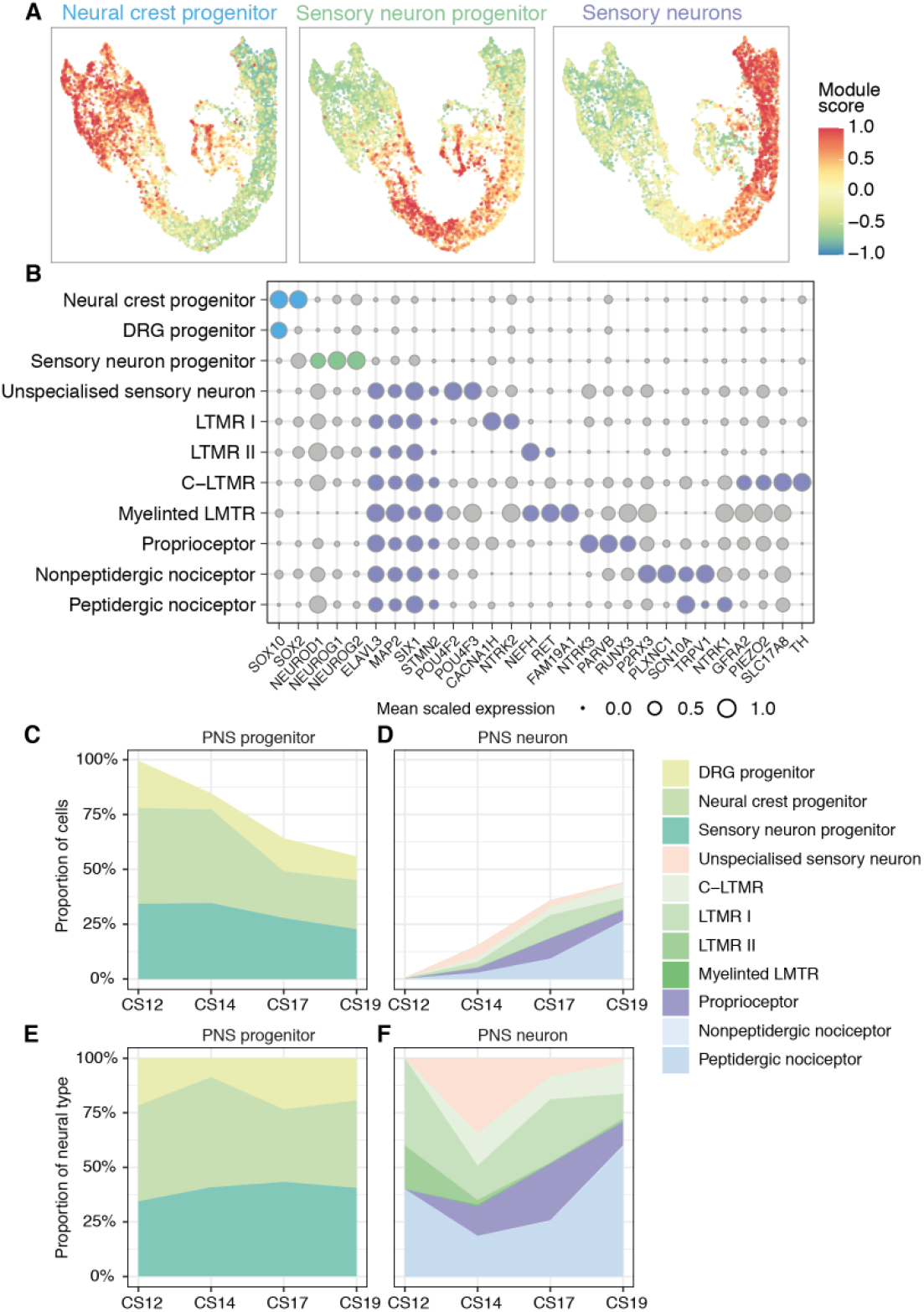
Classification of peripheral nervous system cells in human. **(A)** Gene module scores for progenitors, sensory neuron progenitors, and postmitotic sensory neurons in the peripheral nervous system (PNS) UMAP. **(B)** Bubble plots indicating the expression of markers used to identify PNS cell types. Genes chosen for cell assignment are colored; grey circles correspond to markers not used for the selection of a specific population. Circle size indicates mean scaled gene expression levels. **(C, E)** Fractions of progenitors and **(D, F)** neurons in the PNS during Gestational Week (W) 4 to 7. For (C) and (D), the data are proportional to the total neurons and progenitors at each timepoint. (E) and (F) are the same data within (C) progenitors or (D) neurons as a proportion of the number of neural cells.

We next established a knowledge matrix of dorsal root ganglia expressed genes that distinguish cell types and used this to classify PNS cells into subtypes (FIG 5B). Overall, the classification was similar to mouse, with several transcription factors characteristic of terminal neuronal subtypes in the adult expressed in embryonic sensory neurons (FIG 5B, S5A). Gene expression programmes characteristic of mechanoreceptor (LTMRs), proprioceptor, peptidergic and non-peptidergic neurons were evident. Moreover, the expression profile of genes specific to distinct classes of PNS neurons agreed with recent findings of a single neurogenic trajectory from progenitors through a transcriptionally unspecialized state that then diversifies into distinct sensory lineages (Sharma et al., 2020; Soldatov et al., 2019).

The proportion of neural crest cells and neuronal progenitors decreased from CS12 to CS19 as the proportion of peripheral neurons increased (FIG 5C-F). No PNS neurons were detected at the earliest timepoint (CS12), and a low percentage of terminal neuronal subtypes appeared from CS14 up to CS19 (FIG 5D), indicating a delayed neurogenesis of the PNS system in comparison to the CNS. This suggests that the diversification and specialization of neurons in the PNS occurs at later developmental stages than in the CNS.

### Dynamics of neurogenesis in the human spinal cord

Since the single cell transcriptomic atlas of the mouse neural tube captured temporal changes in progenitor and neuronal populations (Delile et al., 2019), we performed a similar analysis in the human spinal cord. Despite the reduced temporal resolution of the dataset, with four different stages from a period of approximately two weeks, changes in the proportions of progenitors and neurons were evident (FIG 1B,6A,B). The change in the relative proportions of pMN and p3 progenitors were consistent with the measured changes of progenitor domain sizes in the spinal cord over time in human embryos (FIG 6A,B) (Rayon et al., 2020). Extending the analysis to all progenitors and neuronal subtypes revealed that at CS12 ventral progenitors and V1, V2, MN and V3 neurons were more abundant than dorsal subtypes. The relative abundance of neural progenitors decreased over time as neurons increased (FIG 6C,D). Between CS14 and CS19 the generation of neurons in ventral domains appeared to slow (FIG 6D). By contrast, the rate of neurogenesis in dorsal domains increased, with proportionally more dorsal than ventral neurons by CS19 (FIG 6B,D). This is consistent with the dynamics of neuronal subtype differentiation observed in other vertebrates (Kicheva et al., 2014).

**Figure 6.**
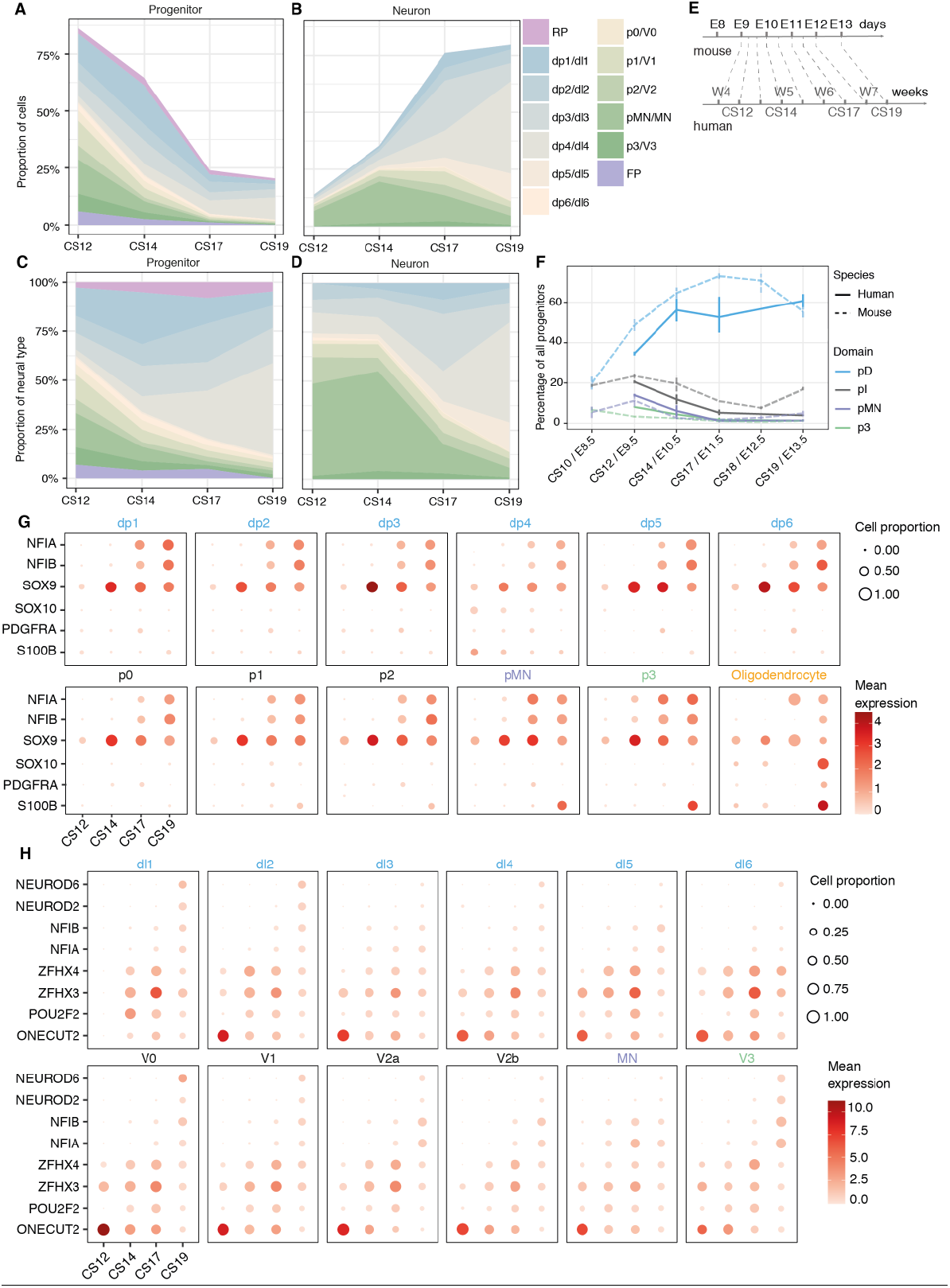
Dynamics of progenitor and neuron proportions in the developing spinal cord. **(A, C)** Proportions of progenitors and **(B, D)** neurons in the spinal cord during W4 to W7. For (A) and (B), the data are proportional to the total neurons and progenitors at each timepoint. (C) and (D) are the same data as a proportion of (C) progenitors or (D) neurons. **(E)** Comparison schematic of stages in mouse and human. **(F)** Comparison of the ratio of progenitors between mouse and human grouped in broad territories: pD (dp1-dp6); pI (p0-p2). Vertical bars indicate the range around the mean of proportions per replicate. **(G)** Expression of gliogenic markers in progenitors and oligodendrocytes. **(H)** Expression of the temporal transcription factor code in neurons. The size of the circles indicates the proportion of cells that express the gene per stage and domain, and the colour indicates the mean expression levels.

We next compared mouse and human spinal cord development by aligning the datasets using major developmental events observed in the two species (Rayon et al., 2020) (FIG 6E). We grouped the 11 neural progenitor domains into dorsal interneuron progenitors (pD; dp1-dp6), intermediate interneuron progenitors (pI; p0-p2), pMNs, and ventral interneuron progenitors p3. A broadly similar pattern of changes in the proportions of cell types were evident in mouse and human, albeit the tempo of human development was considerably slower than mouse. More detailed inspection indicated that at CS12, the proportion of p3 and pMN ventral progenitors compared to pI and pD was higher in human than mouse (FIG 6F). The increased proportion of ventral progenitors persisted at CS14, but by CS17 the size of the ventral domains was comparable between mouse and human (FIG 6F). There was a reduction in the proportion of progenitors in the pD domains from CS14 to CS17, consistent with a slower rate of dorsal neurogenesis in human indicative of a delay in dorsal neurogenesis (FIG 6F). Together, the analysis indicated that, although mouse and human display an overall similar pattern of neurogenesis, the human neural tube has a higher initial proportion of pMN and p3 progenitors and these undergo a higher relative rate of neurogenesis. By contrast, dorsal neurogenesis persists for a longer period in human than mouse.

Finally, we analyzed the sequential expression of gliogenic markers in progenitors. *SOX9* was detected at low levels across progenitor domains from CS12, followed by the detection of *NFIA/B* at CS17 at higher levels in ventral progenitors, its expression increasing in dorsal progenitors by CS19 (FIG 6G). The expression of *SOX9* and *NFIA/B* in progenitors and neurons was consistent with immunohistochemical assays of the embryonic human spinal cord (Betters et al., 2010; Dady et al., 2021; Deneen et al., 2006; Rayon et al., 2020). The expression of NFIA correlates with the onset of gliogenesis and is observed at CS15 in ventral progenitors but delayed until ∼CS18 in dorsal regions (Betters et al., 2010; Dady et al., 2021; Deneen et al., 2006; Rayon et al., 2020). In addition, we identified a small number of cells (n=77), present from CS12 onwards, expressing *SOX10, SOX9, PDGFRA* and *S100*, characteristic of oligodendrocytes (FIG 6G).

### A conserved temporal code for the specification of neuronal identity

In parallel to the spatial patterning of neurons, a temporal transcription factor code has been identified in mouse and is responsible for the further diversification of neuronal identity throughout the central nervous system (Sagner et al., 2020). To investigate if this temporal code is conserved in humans, we examined the expression of the transcription factors that correlate with the early and late born neuronal classes in all dorsoventral domains. Consistent with the analysis of gene expression in the mouse neural tube, human neurons present at the earliest timepoint (CS12) express Onecut-family members but little if any *POU2F2, ZFHX2-4, NFIA/B/X* or *NEUROD2/6*. By CS14 many neurons expressed *POU2F2* and *ZFHX2-4*, characteristic of neurons born at intermediate times in mouse. By contrast, expression of *NEUROD6, NEUROD2, NFIA* and *NFIB*, which identify later generated neurons in mouse, were only detected in neurons by CS19. An exception was MNs, where NFI factors were detected from CS14 (FIG 6H), similar to the earlier expression of these TFs in mouse MNs. The relatively sparse expression of late born neuronal markers in dI3-dI6 in the CS19 sample, suggests that neurogenesis in the human developing spinal cord continues at later timepoints, consistent with an extended neurogenic period in comparison to mouse. Collectively, these results indicate a conserved temporal code for the diversification of neurons that is temporally protracted in the dorsal portion of the spinal cord in human compared to mouse.

### Transcriptional dynamics during neurogenesis in mouse and human

Next, to reconstruct and compare gene expression dynamics during the differentiation of specific neuronal subtypes we used RNA velocity (La Manno et al., 2018) to analyse the temporal dynamics of gene expression. We inferred RNA velocity using scVelo (Bergen et al., 2020) and modelled neurogenic lineages with CellRank (Lange et al., 2020), which infers developmental trajectories from splicing information and transcriptional similarity. For the analysis, we included mouse samples from E9.5 to E13.5 and the CS12, CS14 and CS19 human datasets. Because these methods are very sensitive to the average counts per gene in each dataset, we excluded the CS17 sample, which has lower average gene counts than the other human samples (FIG S6A).

The inference of neurogenic trajectories from the mouse dataset was overall better resolved than in human, likely due to the higher temporal resolution of the mouse datasets compared to the human. Nevertheless, reconstructing the differentiation of neurons of several dorsoventral domains in mouse and human (p3, pMN, p2, p1, dI4, dI3, dI2) showed single convergent neurogenesis paths from progenitors to neurons in each case (FIG 7A, FIG S6B). Moreover, the analysis highlighted a temporal trajectory within some progenitors including p3, pMN, and p2 in mouse (FIG 7A, FIG S6B), which is consistent with the temporal programme of gene expression in progenitors and the gliogenic switch in the spinal cord, where neural progenitors from E11.5 give rise to neurons and glia (Kang et al., 2012; Sagner et al., 2020).

**Figure 7.**
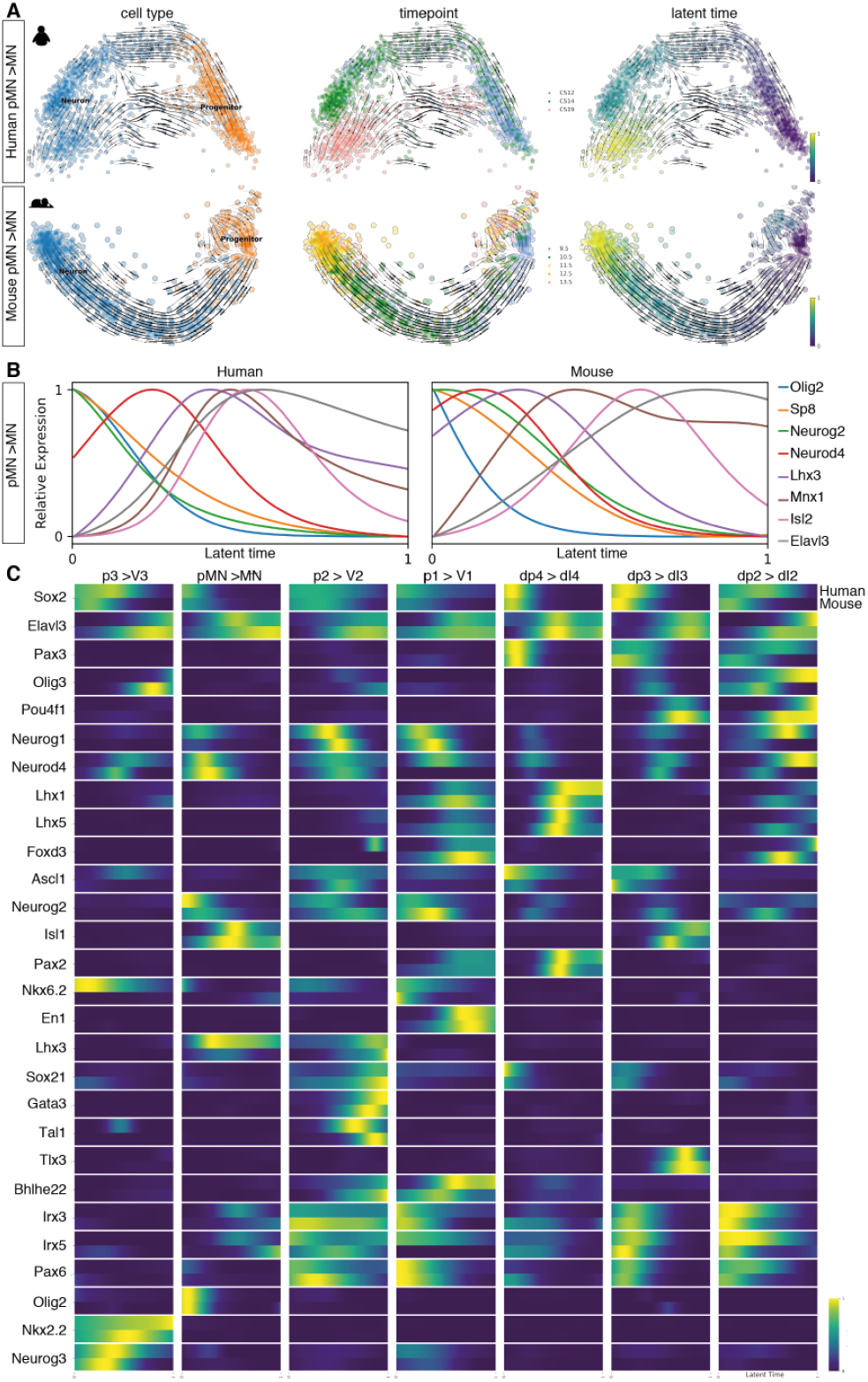
Neurogenic trajectories in the developing spinal cord. **(A)** PCA of the pMN to MN trajectory in human and mouse indicating the RNA-velocity trajectories depicted by black arrows in the PCA. PCAs annotated by cell type on the left, by timepoint in the middle, and by latent time on the right. Earliest latent time points indicated in dark blue and latest in yellow on the latent time color scale. **(B)** Smoothed expression profile of the reconstructed motor neuron differentiation trajectory using the calculated latent times in human and mouse for selected genes. **(C)** Heatmap of the normalised gene expression (blue, low; yellow, high) of genes involved in neurogenesis for each neuronal class as a function of latent time in human (top bar) and mouse (bottom bar).

Latent time estimations of the probability of a cell to reach the differentiated neuronal state allowed directional ordering along the trajectories and the inference of smoothed gene expression dynamics in latent time. In both mouse and human the neurogenic trajectory from pMN to MN confirmed the expected sequential progression of gene expression during differentiation: progenitor genes including Olig2 were expressed earlier in latent time, followed by neurogenic genes such as Neurod4 and finally markers of post mitotic neurons Elavl3 and Mnx1 were detected at the latest latent timepoints (FIG 7B). We then examined the gene expression dynamics for the differentiation of V3, V2, V1, dI4, dI3 and dI2 neurons in mouse and human (FIG 7C, FIG S6C). Heatmaps of the dynamics of gene expression in latent time were generally similar in the two species, and resembled the known dynamics of the genes as well as the pseudotemporal ordering observed in the in scRNAseq in mouse (FIG 7C) (Delile et al., 2019). Overall, these analyses suggested the genetic programmes for the differentiation of neurons in the developing spinal cord are conserved in mammals.

### Co-expression of Olig2 and Nkx2.2 in the developing spinal cord

At early stages of neural development in mouse and chick, pMN cells express OLIG2 but not NKX2.2 and give rise to MNs. This distinguishes pMN progenitors from NKX2.2 expressing p3 cells (Briscoe et al., 1999; Novitch et al., 2001). At later developmental stages, once progenitors finish generating MNs, oligodendrocyte precursor cells (OLPs) in the region of the mouse neural tube previously occupied by pMN cells co-express OLIG2 and NKX2.2 and produce oligodendrocytes (Richardson et al., 2000; Zhou et al., 2001). By contrast, in human embryos a substantial number of OLIG2 and NKX2.2 coexpressing cells are observed in the pMN domain at early developmental stages, when MNs are still being generated ((Marklund et al., 2014); FIG 8A,B, S7A,B). These are thought to represent a pMN subdomain in human (Marklund et al., 2014). To define the molecular identity of OLIG2 and NKX2.2 co-expressing cells, we analysed Sox2+ progenitors that expressed either Nkx2.2 or Olig2 in mouse and human. In mouse, the number of *Olig2+/Nkx2*.*2*+ coexpressing cells comprised less than 5% of the total number of Nkx2.2 or Olig2 expressing cells until E11.5. By contrast, the proportion of OLIG2/NKX2.2 double positive (DP) cells was ∼10% in equivalent staged human embryos (FIG 8C, S7C,D). By E12.5 in mouse, at the onset of gliogenesis, the number of the *Olig2*+/*Nkx2*.*2+* coexpressing progenitors had increased to ∼15%, correlating with the appearance of OLPs in the developing spinal cord. This increase was not detected in human, where the ratio of *OLIG2*+/*NKX2*.*2+* coexpressing progenitors in human decreased at CS17 and CS19 (FIG 6C).

**Figure 8.**
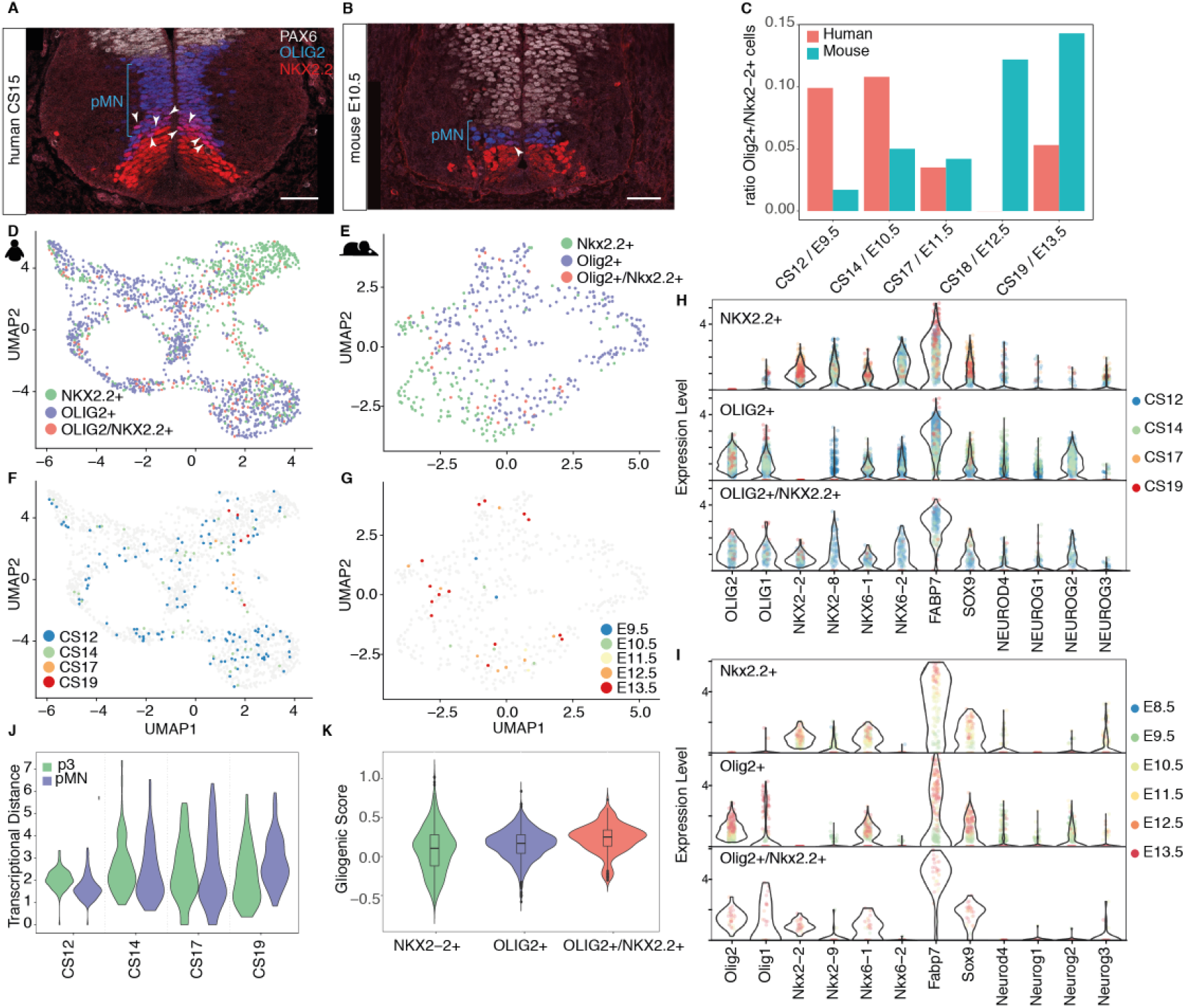
Overlapping expression of OLIG2 and NKX2.2. **(A, B)** Expression of ventral progenitor markers PAX6 (white), OLIG2 (blue), and NKX2.2 (red) in transverse sections of (A) human and (B) mouse cervical neural tube at CS15 and E10.5 respectively. Scale bars, 50μm. **(C)** Ratio of Olig2+/Nkx2.2+ double positive expressing cells within all cells expressing Nkx2.2 or Olig2. **(D-G)** Human UMAP of all cells expressing Nkx2.2 or Olig2 in (D) human (E) and mouse. In (F, G) the double positive progenitors are color coded by timepoint in (F) human and (G) mouse. **(H, I)** Violin plots of selected genes involved in pMN and p3 neurogenesis and progenitor maturation in (H) human and (I) mouse. Cells are labeled by timepoint. **(J)** Transcriptional distance of double positive progenitors to pMN and p3 cells. The closer to 1 the more similar to the population. **(K)** Gliogenic score of cells expressing *OLIG2, NKX2*.*2* or both genes in human.

To investigate whether Olig2+/Nkx2.2+ coexpressing cells had a distinct transcriptional signature, we compared DP cells with Olig2+/Nkx2.2-pMN and Olig2-/Nkx2.2+ p3 progenitors. UMAPs of mouse or human cells showed that DP cells projected to regions of the embedding that contained either pMN or p3 cells (FIG 8D,E). DP cells failed to cluster according to developmental stage (FIG 8F,G). Moreover, analysis of pMN and p3 markers in human and mouse DP cells showed a similar transcriptional signature, which closely resembled a combination of the pMN and p3 signatures (FIG 8H,I). To determine the similarity of DP cells to either pMN cells or p3 cells, we averaged the expression profiles for each cell type per time point and calculated the distance between DP cells and pMN or p3 cells in human. This suggested that double positive cells were somewhat closer in gene expression space to pMN cells at CS12 and CS14 than to p3 cells (FIG 8J), but the degree of similarity with p3 cells increased at later timepoints (FIG 8J). Together, this analysis suggests that human *OLIG2+/NKX2*.*2+* coexpressing progenitors resemble an amalgamation of pMN and p3 identities.

The co-expression of OLIG2 and NKX2.2 in mouse and chick is indicative of OLPs. For this reason, the DP cells in human have been proposed to be OLPs (Marklund et al., 2014). Consistent with this, human DP cells expressed *FABP7, OLIG1* and *OLIG2*, characteristic of gliogenic progenitors, similar to mouse DP cells from older embryos (FIG 8H, I). Whereas the expression of gliogenic markers in DP cells from mouse at early timepoints was low, the levels of *FABP7, OLIG1* and *OLIG2* in human double positive progenitors were consistently high across timepoints (FIG 8H). Likewise, we observed an increased gliogenic score in the double positive progenitors compared to pMN and p3 progenitor populations (FIG 8K). Taken together, these results are consistent with DP cells representing OLPs. This would indicate that OLPs are more abundant at earlier developmental stages in the human neural tube than at the equivalent mouse stages, representing another example of heterochronic differences between rodents and primates. In chick and mouse, OLPs appear to arise mainly after pMN progenitors have ended MN generation, but the lineage relationship and the neurogenic or gliogenic potential of individual cells needs clarification (Leber and Sanes, 1995; Leber et al., 1990; Park et al., 2002; Richardson et al., 2000; Wu et al., 2004). In zebrafish, MNs and oligodendrocytes have been proposed to arise from distinct cell lineages that initiate *Olig2* expression at different times (Ravanelli and Appel, 2015). It will be important to establish the lineage relationships between pMNs, MNs and OLPs in human, and establish the neurogenic or gliogenic potential of double positive cells.

### Online access to mouse and human neural tube transcriptome data

The availability of mouse and human transcriptome data from the developing neural tube and dorsal root ganglia provides a resource for developmental biologists studying the embryonic nervous system and for stem cell biologists aiming to refine differentiation protocols for the generation of specific neuronal subtypes. The *in vivo* characterization of cell diversity in mouse and human will help identify the neuronal subtypes involved in locomotor circuits and provides a foundation to investigate subtype-specific and species-specific features that control connectivity and function. To provide access to both the human and mouse transcriptome data we have established an interactive web application (https://shiny.crick.ac.uk/scviewer/neuraltube/). This will allow rapid and easy investigation of the expression of specific genes in all datasets described in this analysis (see Movie S1).

## Materials and Methods

### Sample Processing

Human embryonic material (GW 4-7) was obtained from the MRC/Wellcome-Trust (grant #006237/1) funded Human Developmental Biology Resource (HDBR57, http://www.hdbr.org) with appropriate maternal written consent and approval from the London Fulham Research Ethics Committee (18/LO/0822) and the Newcastle and North Tyneside NHS Health Authority Joint Ethics Committee (08/H0906/21+5). HDBR is regulated by the UK Human Tissue Authority (HTA; www.hta.gov.uk) and operates in accordance with the relevant HTA Codes of Practice. Embryos were freshly collected on Hibernate-E medium (A12486, Gibco) supplemented with 2% B-27 (17504001 ThermoFisher Scientific) and 2.5 mL/L Glutamax (3505006, Gibco) and kept on ice until further dissection.

Trunk samples of single embryos were dissected as in (Delile et al., 2019). Briefly, samples were placed in Hanks Balanced Solution without calcium and magnesium (HBSS, 14185045, Life Technologies) supplemented with 5% heat-inactivated foetal bovine serum (FBS). For neural tube and dorsal root ganglia dissection, the superficial layers of skin, limbs and remaining organs were removed, whereas the somites, cartilage primordium, dorsal root ganglia and neural tube were left intact. Dissected samples were snipped into smaller pieces and incubated on a FACSmax (Amsbio, T200100) cell dissociation solution containing 10X Papain (30 U/mg, Sigma-Aldrich, 10108014001) for 11 min at 37°C to dissociate the cells and transferred to a HBSS solution with 5% FBS, Rock inhibitor (10 μM, Stemcell Technologies, Y-27632) and 1X of non-essential aminoacids (Thermo Fisher Scientific, 11140035). Single cells were disaggregated through pipetting and filtered, and quality control was assayed by measuring live cells versus cell death, cell size and number of clumps on an EVE cell counter. Each sample was prepared at a concentration of 600-1700 cells/µl and viability ranged between 70-96%. Samples with a viability above 70% were used for sequencing.

### Single-cell RNA sequencing

Single cell suspensions were loaded for each sample into a separate channel of a Chromium Chip G for use in the 10X Chromium Controller (cat: PN-1000120). The cells were partitioned into nanolitre scale Gel Beads in emulsions (GEMs) and lysed using the 10x Genomics Single Cell 3′ Chip V3.1 GEM, Library and Gel Bead Kit (cat: PN-1000121). The v2 kit was used for the E8.5 mouse as in (Delile et al., 2019). cDNA synthesis and library construction were performed as per the manufacturer’s instructions. Libraries were prepared from 10µl of the cDNA and 12 cycles of amplification. Each library was prepared using Single Index Kit T Set A (cat: PN-1000213) and sequenced on the HiSeq4000 system (Illumina) using the configuration 28-8-98 on a single-index-paired-end run. Libraries were sequenced on independent flow cells for each sample and split across multiple lanes.

### Quality control and filtering

Gene expression was quantified from FastQ files using Cell Ranger (3.1.0) and the GRCh38-3.0.0 and mm10-3.0.0 indexes for human and mouse respectively. A counts matrix was generated for each library using Cell Ranger count, submitting all technical replicates of a library together. Putative cells reported by Cell Ranger in the filtered output matrix were inspected manually to remove droplets that reported an extremely high or low number of genes or UMI or had a high proportion of mitochondrial UMI. The table of thresholds applied to each replicate can be found in Table S1.

### Cell-cycle and gene module scoring

Each cell was assigned a score based on the expression of a pre-determined list of cell-cycle gene markers (cc.genes, supplied with Seurat) with the CellCycleScoring function implemented in Seurat (Stuart et al., 2019). Mouse gene names were converted from human using biomaRt Ensembl-release 93 gene annotations. Additionally, gene module scores were calculated using the AddModuleScore function in Seurat for each gene module described in Table S2. Gene module scores were used for visualisation purposes on reduced dimension plots and to identify clusters of cells that were likely non neural cells that were removed from further analysis. The gliogenic score of *OLIG2*+/*NKX2*.*2*+ double positive cells in human was determined using Seurat’s AddModuleScore function with a set of known gliogenic markers (*FABP7, SOX9, SOX10, PDGFRA, CSPG4, FGFR3, FGFBP3, DBI, SLC1A3, HOPX, ALDH1L1*).

### Normalisation, integration and dimension reduction

Replicate datasets were subsequently filtered to remove contaminant cells (such as mesoderm and blood). The resulting datasets were then normalised and integrated using the SCTransform/IntegrateData workflow implemented in Seurat 3.2.2 (Stuart et al., 2019). For SCTransform, 3000 variable features were selected and cell cycle effects regressed using the difference in cell cycle scores (S-G2M). Replicates were then integrated together to create the time point and species datasets.

Principal components of integrated datasets were calculated using RunPCA with 3000 variable features to provide the first 70 components. UMAP and tSNE reductions were calculated using the RunUMAP and RunTSNA using the first 40 principal components.

### Cell type classification

We classified cells using Antler (https://juliendelile.github.io/Antler/) as described in (Delile et al., 2019). Briefly, cells were classified based on the expression of known marker genes in a two-step process that first identified broad cell types before classifying neurons and progenitors further into 12 and 13 dorsoventral sub-types respectively. The knowledge matrices (Table S2 and S3) were adapted from (Delile et al., 2019), converting mouse to human gene identifiers and replacing *Tubb3* with *STMN2* and *MAP2*, as *TUBB3* levels were low in human. In the human, additional marker genes were added for the PNS classification and for the identification of oligodendrocytes (Table S2).

### Mouse and human correlation analysis

First, a set of orthologous genes with a 1:1 correspondence were sought between human and mouse using Ensembl biomaRt and release-93 annotations. Gene sets were then defined. The ‘transcription factor’ gene set was defined as the unique set of genes with which the GO term ‘GO:0003700’ was associated, using human biomaRt connection.

Expression matrices were subset by cell type and gene set. Human gene names were converted to mouse gene names and genes with no 1:1 homologue were discarded. Representative expression profiles per cell type were calculated using the mean gene expression. Each representative profile was correlated using the ‘cor.test’ function in R and the Pearson method.

### Comparison to bulk in vitro RNA-seq data

Data from ventral (p3, pMN, p2, p1, p0) domain progenitors and neurons were aggregated and averaged per timepoint. A subset of 1448 highly expressed genes with varying dynamics along the neurogenic differentiation trajectory was chosen to compare this pseudo-bulk data with bulk *in vitro* data published previously (Rayon et al., 2020). Data from each timepoint was normalised between 0 and 1 to improve comparability and Spearman’s correlation coefficient across all average expressions was calculated between each *in vivo* and *in vitro* timepoint.

### Correlation analysis for *PAX7*

To investigate PAX7 expression in floorplate (FP) cells, Spearman’s correlation coefficient was used to select the 30 genes that have the strongest positive correlation with PAX7 in cells annotated as floorplate. The table with the mean expression levels of the genes in human FP cells, pD cells and mouse FP can be found on Table S4.

### RNA Velocity

Spliced/unspliced count matrices were generated from aligned reads using the Velocyto (La Manno et al., 2018) command line interface tool with the run10x command and default arguments. For reference genomes and repeat masks, GRCh38-3.0.0 and mm10-3.0.0 were used for human and mouse respectively.

Only cells with Antler classifications were used for downstream analysis without the SCTransform/IntegrateData workflow implemented in Seurat 3.2.2 normalization step (Stuart et al., 2019). It was noted that CS17 data had lower counts per gene than other timepoints and this produced artefacts in normalisation and scaling pre-processing steps for PCA (Figure S6A). As a result, CS17 was not included in trajectory analysis. Mouse E8.5 data was also excluded as it had no corresponding timepoint in human data. Initial inspection of p0, dl6, dl5 and dl1 domains revealed insufficient number of cells to model the progenitor-to-neuron transition and so these domains were not included in further trajectory analysis.

For downstream analysis, ScanPy was used to normalise and log transform the data. To subset genes, 2000 highly variable genes were selected for each timepoint separately, and genes common to all timepoints were retained. This was done to reduce batch effects between timepoints. To these highly variable genes, a curated list of 177 genes known to be involved in neural tube development was added. Gene selection, PCA and velocity modelling were performed on each domain independently. ScVelo (Bergen et al., 2020) was used to recover ‘latent time’ dynamics, infer velocities on ‘stochastic’ mode, and embed these velocities in PCA space.

### CellRank Trajectory Analysis

CellRank (Lange et al., 2020) was used to construct neurogenic gene trends. CellRank models cellular dynamics as a Markov chain where transition probabilities are calculated using both RNA velocity and transcriptomic similarity. First, each domain’s terminal states were inferred with the number of terminal states fixed to either two or one depending on whether a bifurcation of progenitor fates was visible in the first two PCA components (for both mouse and human: two terminal states for p3, pMN, p2, p1 and dl2, one for dl4 and dl3). The weight_connectivities parameter, which determines the relative importance of velocity and transcriptomic similarity for terminal state calculation, was adjusted for each domain to achieve a realistic prediction (for p3, pMN, p2, p1, dl4, dl3, dl2: 0.2, 0.7, 0.7, 0.2, 0.7, 0.9, 0.5 respectively in human and 0.2, 0.2, 0.3, 0.2, 0.7, 0.7, 0.5 respectively in mouse). To produce gene trends, a generalised additive model (GAM) was fitted to each gene’s expression pattern through latent time using cellrank’s ul.models.GAM function. This was done with 5 knots and a spline order of 2. This function weights a cell’s contribution to the model by its lineage probability, so progenitors committing to a gliogenic/late progenitor identity did not contribute to neurogenic gene trends. If this GAM failed to fit to a gene’s expression profile, or the GAM’s confidence interval achieved a range larger than 10, the expression profile was fixed to zero.

### Immunostaining and microscopy

Immunohistochemistry on human and mouse spinal cord tissues, and on mouse and human cells was performed as described previously (Rayon et al., 2020). Rabbit anti-PAX6 (Covance PRB-278P-100, 1:500), goat-anti OLIG2 (R&D AF2418, 1:800), mouse anti-NKX2.2 (BD Pharmigen 74.5A5,1:500), mouse anti-NKX6.1 (DHSB F55A10, 1:100), and guinea pig anti-NKX6.2 ((Vallstedt et al., 2001), 1:8000) were used. The anti-NKX6.2 antibody raised against the 11 amino acid N-terminal mouse epitope contained a mismatch (Alanine (A) > Threonine (T)) in the human sequence and did not detect NKX6.2 in human samples.

Cryosections were imaged using a Leica SP8 confocal microscope equipped with a 20x NA 0.75 dry objective, or a Leica SP5 confocal microscope. Z stacks were acquired and represented as maximum intensity projections using ImageJ software. Pixel intensities were adjusted across the entire image in Fiji. The same settings were applied to all images.

### RNAScope FISH with immunohistochemistry

Transverse cryosections (thickness: 14μm) were onto Superfrost PlusTM slides (Thermo ScientificTM 10149870) as in (Rayon et al., 2020). Slides were stored at −80°C until ready to be processed. The RNAscope multiplex fluorescent v2 kit was used per the manufacturer’s instructions for fixed frozen samples (ACD Bio), with the protease treatment step shortened to 15 minutes. The RNAscope Probe-Hs-NKX6-2 (Cat No. 48674) was used. At the end of the RNAscope protocol, sections were fixed in 4% paraformaldehyde for 15 minutes at room temperature and then washed twice in 1X PBS for 5 minutes. Sections were incubated in blocking solution (1% BSA, 0.1% triton-x 100 in 1X PBS) for 30 minutes at room temperature and then incubated in primary antibody in blocking solution (F55A10 DHSB, 1:100) 1h at 4°C. Sections were then washed 3 times for 5 minutes each in 1X PBS, incubated with secondary antibody (donkey anti-mouse Alexa Fluor 488 conjugate, 1:1000) for 30 minutes at room temperature, rinsed in 1X PBS 3 times for 5 minutes each, and mounted with ProLong Gold with DAPI mounting medium (ThermoFisher Scientific). Sections were imaged using a 40X oil immersion lens on a Leica SP8 confocal microscope. Images were taken with the pinhole set to 2AU.

### Data Availability

Single cell RNA sequencing data have been deposited in NCBI-GEO under accession number GSE171892. The R analysis script developed for this paper is available at https://github.com/briscoelab/human_single_cell. Processed single-cell sequencing is available for exploration at our cell browser https://shiny.crick.ac.uk/scviewer/neuraltube/.

## Supplementary information

Table S1. Cell numbers and thresholds applied for each human sample.

Table S2. Knowledge matrix used to identify cell types in human.

Table S3. Knowledge matrix used to identify cell types in mouse (Delile et al., 2019).

Table S4. Mean expression levels of genes correlating with PAX7 expression in human FP, pD and mouse FP.

Movie S1. An interactive web application of the human and mouse transcriptome data.

## Acknowledgements

We are grateful for the human embryonic material provided by MRC/Wellcome Trust (MR/R006237/1) Human Developmental Biology Resource and the generous donors whose contributions have enabled this research. We thank past and current members of the Briscoe lab for feedback. We acknowledge the Crick Science and Technology platforms, especially Advanced Sequencing, Scientific Computing, Bioinformatics and Biostatistics group, Mary Green from Experimental Histopathology Laboratory, and Marc Pollitt from Scientific Computing. Manuela Melchionda provided the embryo sections; Katherine Exelby designed the spinal cord diagram. We thank Despina Stamataki, Andreas Sagner, M. Joaquina Delás, Tiago Rito and Tom Frith for comments on the manuscript. The authors are supported by the Francis Crick Institute, which receives its core funding from Cancer Research UK, the UK Medical Research Council, and the Wellcome Trust (all under FC001051). T.R. and. J.B. are also supported by the Wellcome Trust (215116/Z/18/Z). T.R. received funding from an EMBO long-term fellowship (ALTF 328-2015). This research was funded in whole, or in part, by the Wellcome Trust (FC001051). For the purpose of Open Access, the author has applied a CC BY public copyright licence to any Author Accepted Manuscript version arising from this submission.

## Author contributions

Conceptualization: T.R., J.B.; Methodology: T.R., R.M., C.B., J.B.; Software: R.M., C.B.; Formal analysis: R.M., C.B; Investigation: T.R., J.B.; Writing - original draft: T.R., J.B.; Writing - review & editing: T.R., R.M., C.B., J.B.; Supervision: T.R., J.B.; Project administration: T.R., J.B.; Funding acquisition: J.B.

## Supplemental Figure Legends

**Figure S1.**
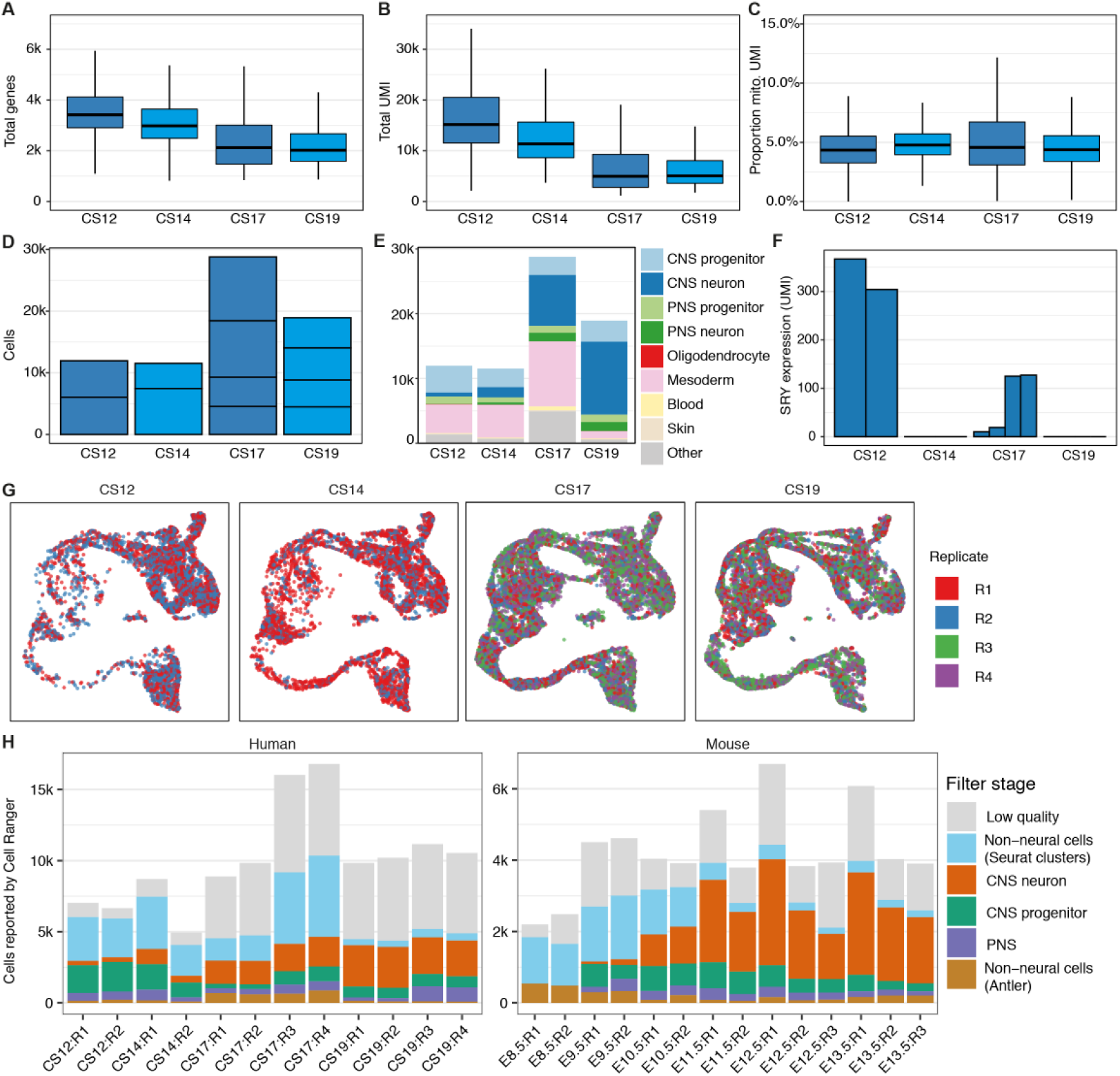
Characterization of the single cell transcriptome dataset. **(A)** Number of genes per cell for each stage. **(B)** Number of unique molecular identifiers (UMI) per cell for each stage. **(C)** Proportion of mitochondria UMI per cell for each stage. **(D)** Number of cells per replicate in each timepoint. **(E)** Number of cells classified per cell type in each stage. **(F)** Expression of the male-specific gene SRY per embryo. **(G)** UMAP of neural cells by timepoint and replicate. **(H)** Number of cells identified by Cell Ranger for each embryo. Low quality cells were removed using (A-C). Curated cells were classified with Seurat and Antler.

**Figure S2.**
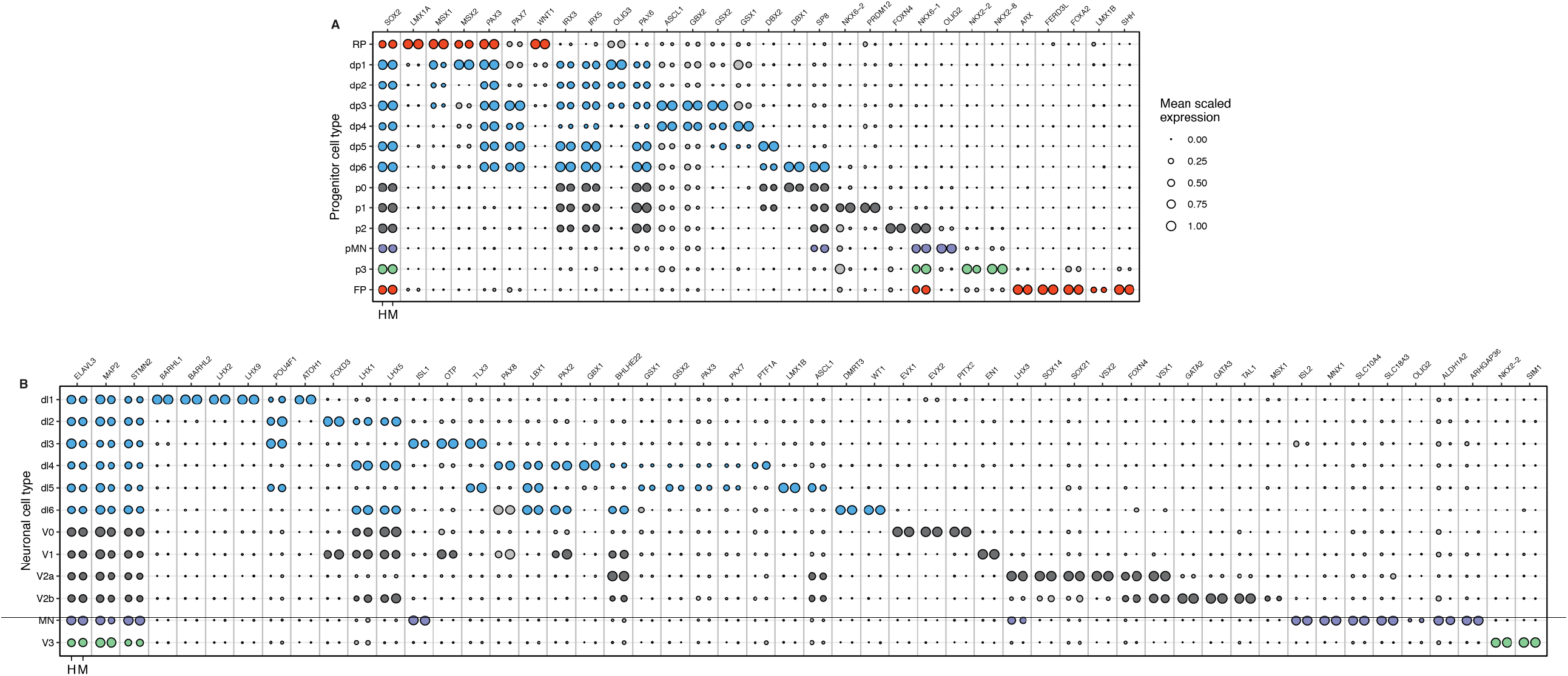
Classification of dorsoventral progenitors and neuronal classes in human and mouse. **(A, B)** Combined bubble plot that depicts the expression of markers used to identify DV domains in human and mouse (A) progenitors and (B) neurons. Genes chosen for cell assignment are coloured; grey circles correspond to markers not used for the selection of a specific population. Circle size indicates mean scaled gene expression levels.

**Figure S3.**
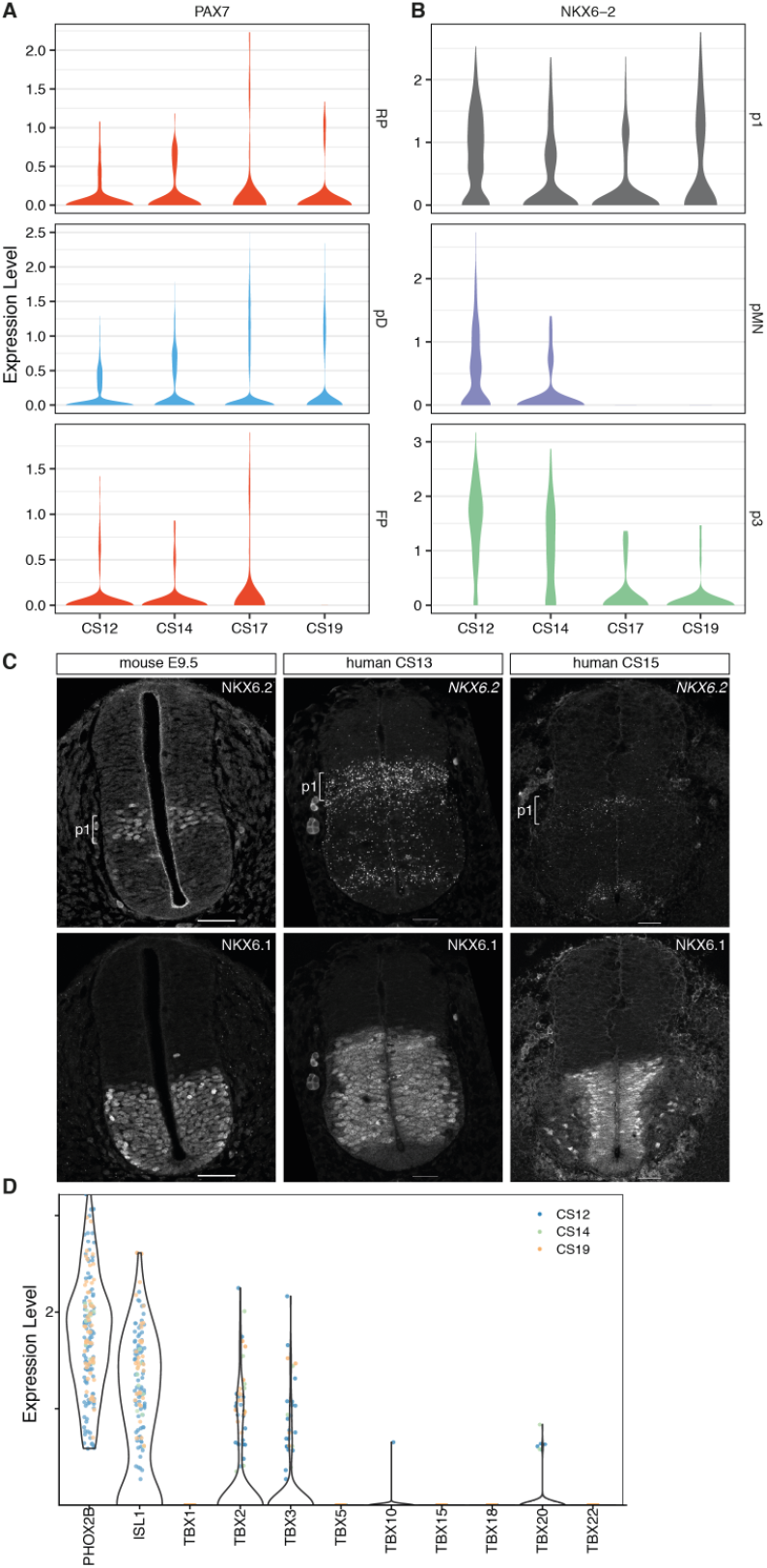
Further characterization of human-specific features of neural progenitors and visceral motor neurons. **(A)** Violin plots of *PAX7* expression in the floor plate (FP), pD (dp1-dp6) and roof plate (RP) at the indicated timepoints. **(B)** Violin plots of *NKX6*.2 expression in p1, pMN and p3 progenitors. **(C)** NKX6.2 and NKX6.1 expression in transverse sections of the mouse and human neural tube at shoulder levels in mouse E9.5, and human CS13 and CS15 embryos. Scale bars, 50μm. **(D)** Violin plots for the expression of the indicated TBX genes in visceral motor neurons classified according to the expression of *PHOX2B* in human color coded by timepoint.

**Figure S4.**
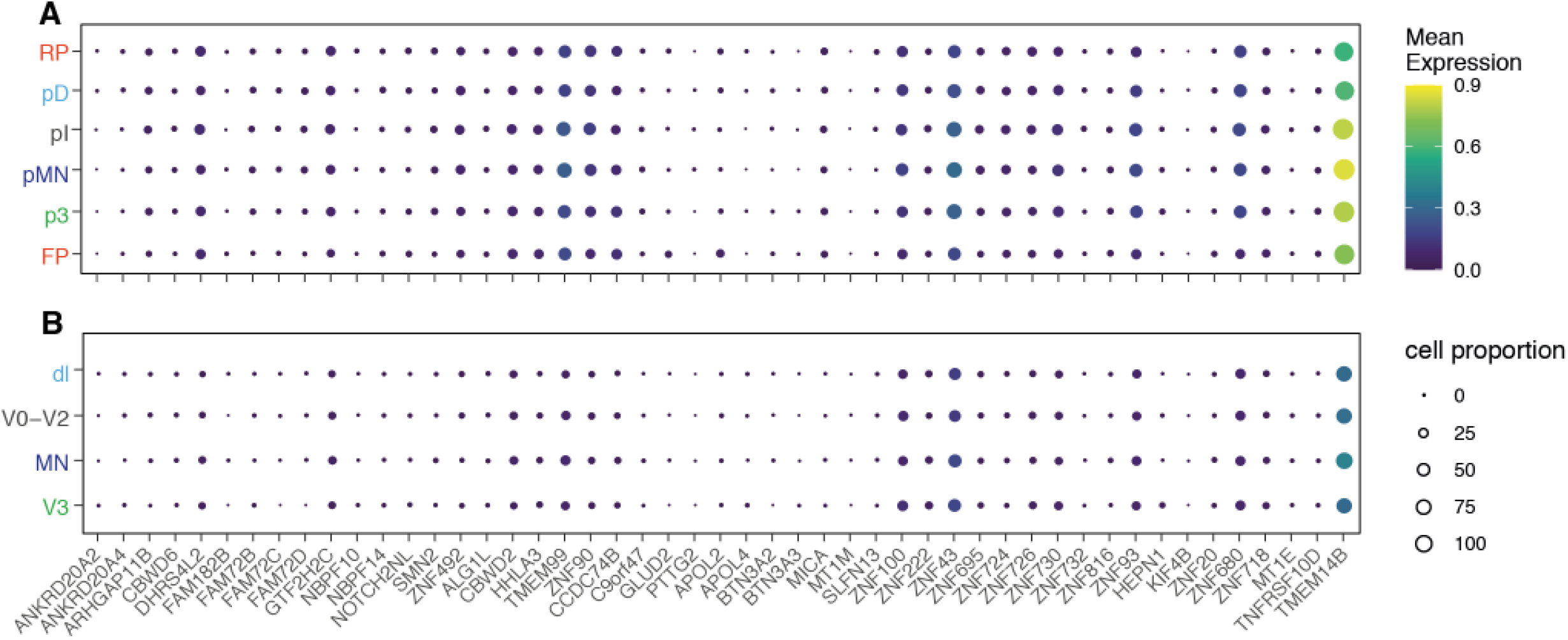
Expression of primate-specific genes in progenitors and neurons of the human developing spinal cord. **(A, B)** Bubble plots that depict the expression of primate specific genes in (A) DV progenitors or (B) neurons. The size of the circles indicates the proportion of cells that express the gene per stage and domain, and the colour indicates the mean expression levels. pD (dp1-dp6), pI (p0-p2), dI (dI1-dI6).

**Figure S5.**
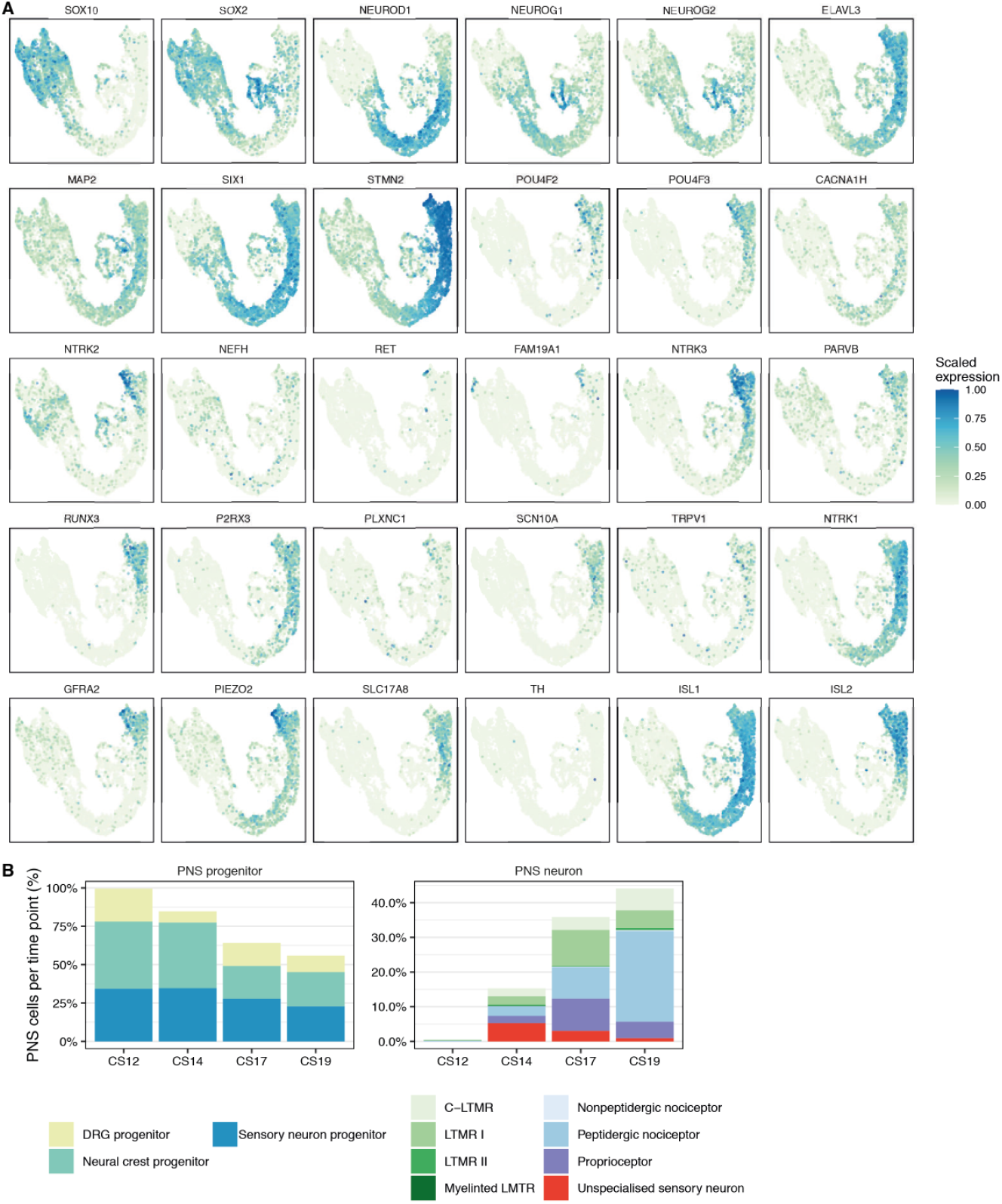
Further classification of peripheral nervous system cells in the developing human embryo. **(A)** Feature plots of genes selected for the classification. **(B)** Proportion of peripheral nervous system (PNS) progenitors and neurons per time point.

**Figure S6.**
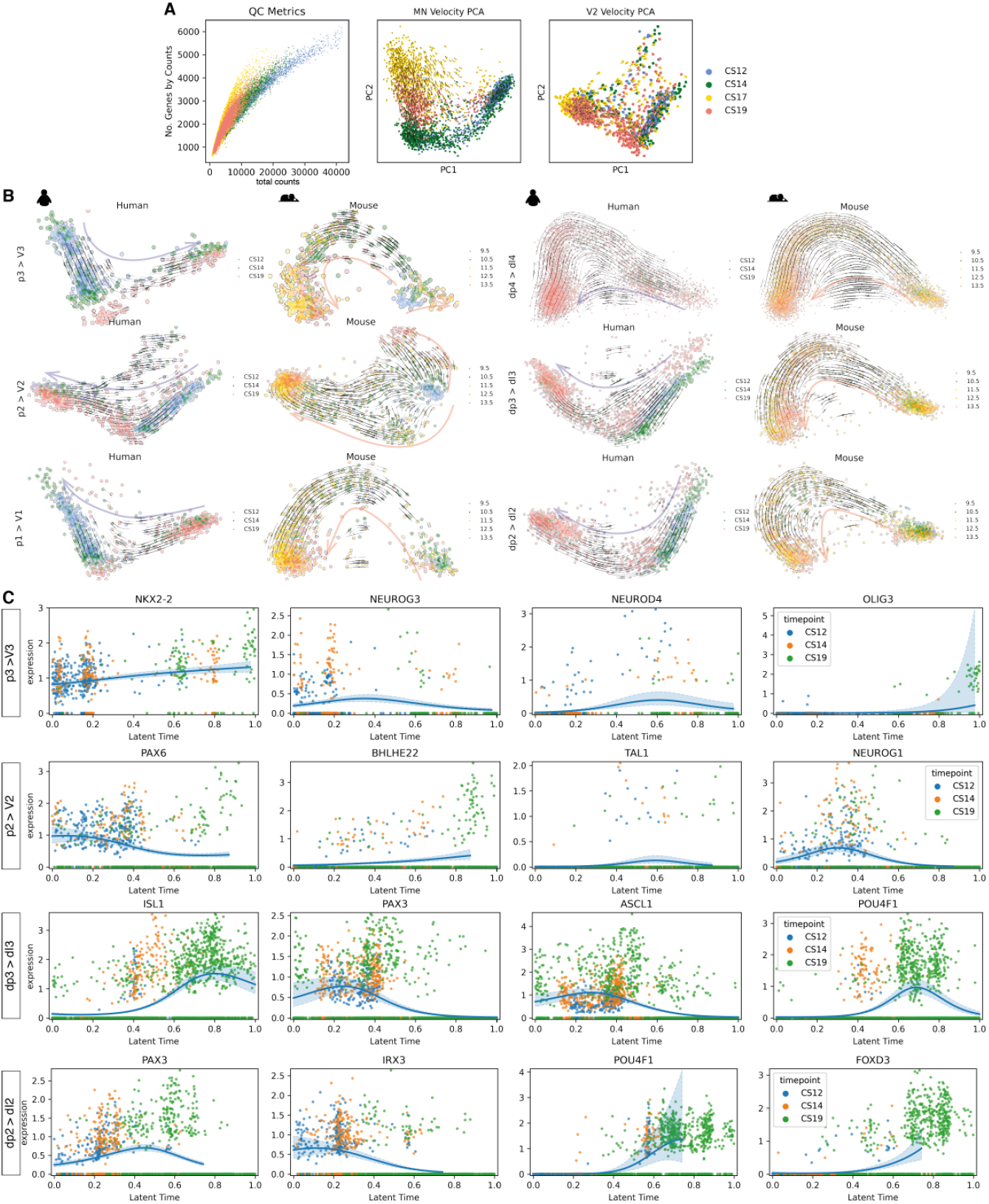
Neurogenic trajectory interference across dorsoventral domains of the developing spinal cord. **(A)** Total counts per number of genes in each timepoint for velocity analysis and PCA of the pMN to MN trajectory in human (middle) and the PCA of the V2 trajectory including the CS17 sample. **(B)** PCA of neurogenic trajectories in human and mouse indicating the RNA-velocity trajectories depicted by black arrows in the PCA for dI2, dI3, dI4, V1, V2 and V3 neurons. PCAs colored by timepoint. Light arrows indicate the inferred trajectory paths. **(C)** Gene expression levels of selected genes per cell along the estimated latent time with the interpolated smoothed expression in human. Cells colored by timepoint.

**Figure S7.**
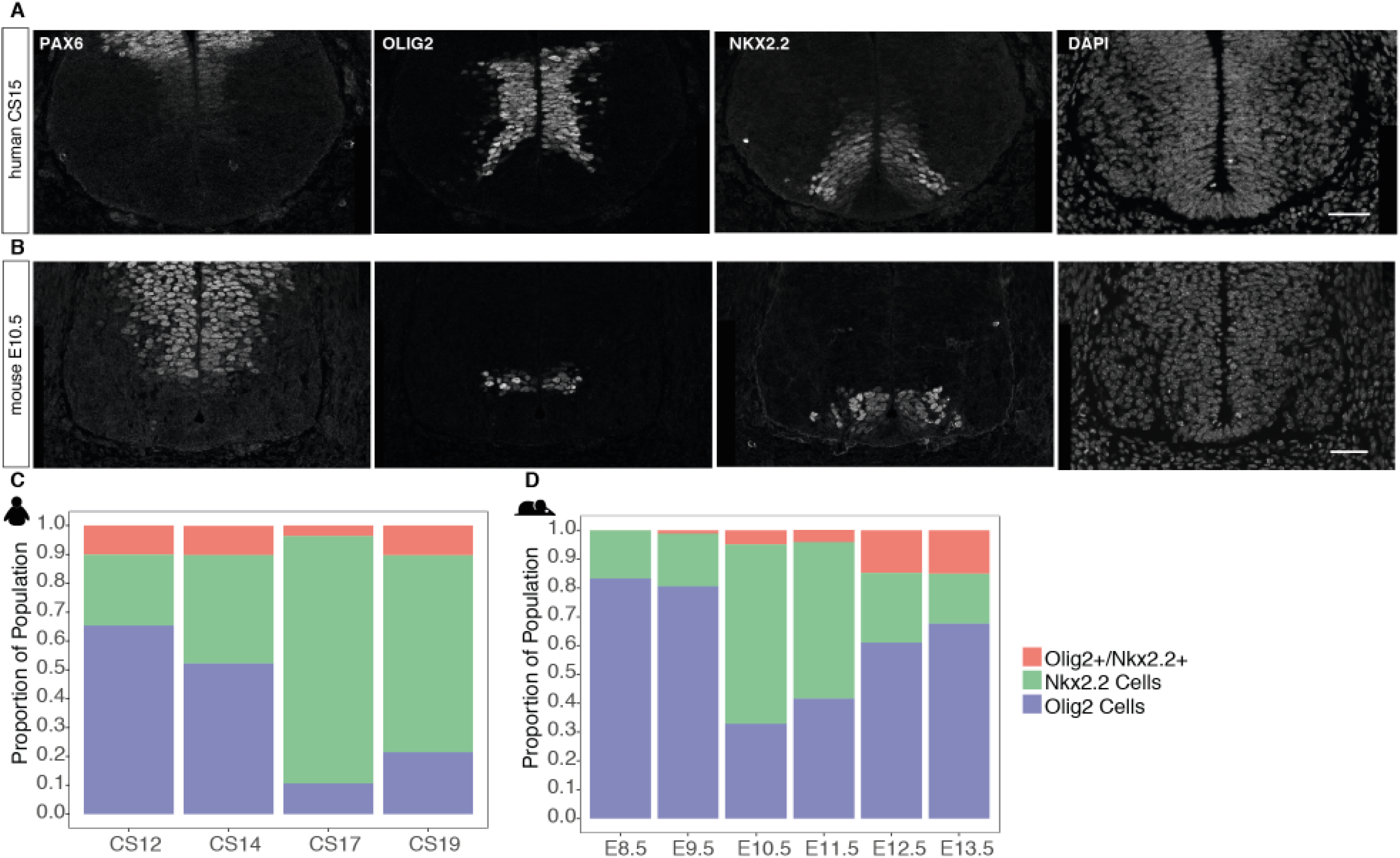
Expression of Olig2 and Nkx2.2 in mouse and human. **(A, B)** Expression of ventral progenitor markers PAX6, OLIG2, and NKX2.2 and DAPI in transverse sections of (A) human and (B) mouse cervical neural tube at CS15 and E10.5 respectively. Scale bars, 50μm. **(C, D)** Proportion of Olig2, Nkx2.2 and co-expressing cells across timepoints in (C) human and (D) mouse.

